# Universal Geometry of Compositional Construction in Prefrontal Cortex

**DOI:** 10.64898/2026.04.23.720375

**Authors:** Maxim Manakov, Mikhail Proskurin, Hanqing Wang, Elena Kuleshova, Andy Lustig, Reza Behnam, Shaul Druckmann, D. Gowanlock R. Tervo, Sue Ann Koay, Alla Y. Karpova

## Abstract

Compositional generation underlies the systematic and essentially unlimited construction of complex concepts from simpler parts, as is foundational to intelligent behavior, but its underlying neural mechanisms remain unclear. Here we reveal a neural implementation of hierarchical compositional construction of abstract sequences. We demonstrate that in an open-ended setting with very sparse feedback, rats innately utilize hierarchical composition to construct adaptive action sequences that would have been difficult to discover from scratch. Prefrontal neural population representations of these abstract sequences adhere to a low-dimensional format that encodes the orderly progression of elemental units comprising the sequence while converging to a sequence-general endpoint. Higher-level compositions in the hierarchy are systematically related to their lower-level constituent parts, reusing much of the representation, while providing context separation and satisfying format constraints. These neural representations are geometrically identical across animals, pointing to a convergent solution for how knowledge is hierarchically assembled via a compositional mechanism.

## Introduction

The ability to systematically construct new, complex behaviors or ideas from simpler parts is a hallmark of intelligence. A longstanding proposal is that composition provides a simple and general mechanism for constructing new knowledge by repeatedly combining concepts in novel ways, making “infinite use of finite means”^1^. A classic example of a compositional system is human language, where a small set of phonemes can be arranged into a much larger set of words that themselves can be arranged into an even larger set of possible sentences^1^. In human cognition, composition is also thought to be at the core of the generative capacity of nonlinguistic thought and reasoning^2–6^. Across animal species, compositional generation is thought to underlie the astonishing variety and complexity of adaptive behaviors that individuals can rapidly discover^7^.

Inherent in the efficacy of compositional mechanisms in enabling complex behaviors is that they extend *beyond* combining the elemental behavioral units (“first-order” composition). Indeed, compositional construction is fundamentally *hierarchical*: once a behaviorally meaningful first-order composite unit has been discovered, it can function as a new, more complex input into subsequent “second-order” composition (and so forth). In this respect, the *efficiency* of composition arises because previously learned units can be flexibly reused in new combinations, but the *power* of composition arises because the behavioral “tool kit” of useful compositional parts progressively expands and becomes more sophisticated, enabling the generation of complex adaptive behaviors that would have been prohibitively difficult to discover from scratch. Recent advances have begun to elucidate how biological^8–10^ and artificial^11^ neural networks may flexibly put together behavioral/computational elements to represent new combinations. While these efforts have focused on signatures of reuse, hierarchical composition requires representations of composite units that go beyond the juxtaposition of their constituent parts. From a mechanistic perspective, in order for composite units to act as inputs to second-order compositional construction, they must also acquire characteristics that identify them as valid “parts”. A central challenge is thus to design experimental settings where both first- and second-order compositions can be observed within the same behavioral framework, permitting us to identify the nature of these characteristics and neural mechanisms that confer them onto constructed composites.

Here we extended our unguided sequence discovery framework^12^, in which rats learn to discover abstract sequences of two elemental action-units (“L” and “R”) matching experimenter-defined rewarded (“target”) patterns, given no external feedback beyond sparse and delayed rewards (Figure 1A). Having previously shown that rats consistently discover sequences that match short target patterns (e.g. “LLR”, “RRLR”, …), we now demonstrate that these behavioral sequences bear hallmarks of first-order composition; and furthermore, rats reuse these learned composite units in second-order composition to match substantially longer, difficult-to-discover targets like “LLRRRLR”. Leveraging this framework, we uncover the neural underpinnings of hierarchical sequence construction by performing cellular-resolution recordings from the anterior medial frontal cortex (amFC) — a region previously implicated in rats in the sequence discovery task^12,13^, and in humans in generative composition^14,15^. Strikingly, we find that the geometry of the neural representations — viewed from a neural population perspective — is quantitatively uniform across animals for both first-and second-order behavioral composition. At a finer-grained level, the neural representations of all compositions conform to a simple format, encoding the orderly progression of “L” and “R” elements toward a sequence-general endpoint. Moreover, consistent with compositional reuse, the representations of second-order compositions closely follow those of their constituent parts. Nevertheless, the representations of second order compositions remain clearly distinct from a simple concatenation of those lower-level representations, thus providing a contextual signal for when a part is reused in a subsequent composition. Crucially, this reorganization is not random but rather constrained to maintain a shared representational format across levels of compositional hierarchy. Our findings thus reveal a novel and convergent representational mechanism in the brain that underlies compositional construction of action sequences, and points to how knowledge can be progressively built upon and flexibly adapted to support intelligent behavior.

**Figure 1.**
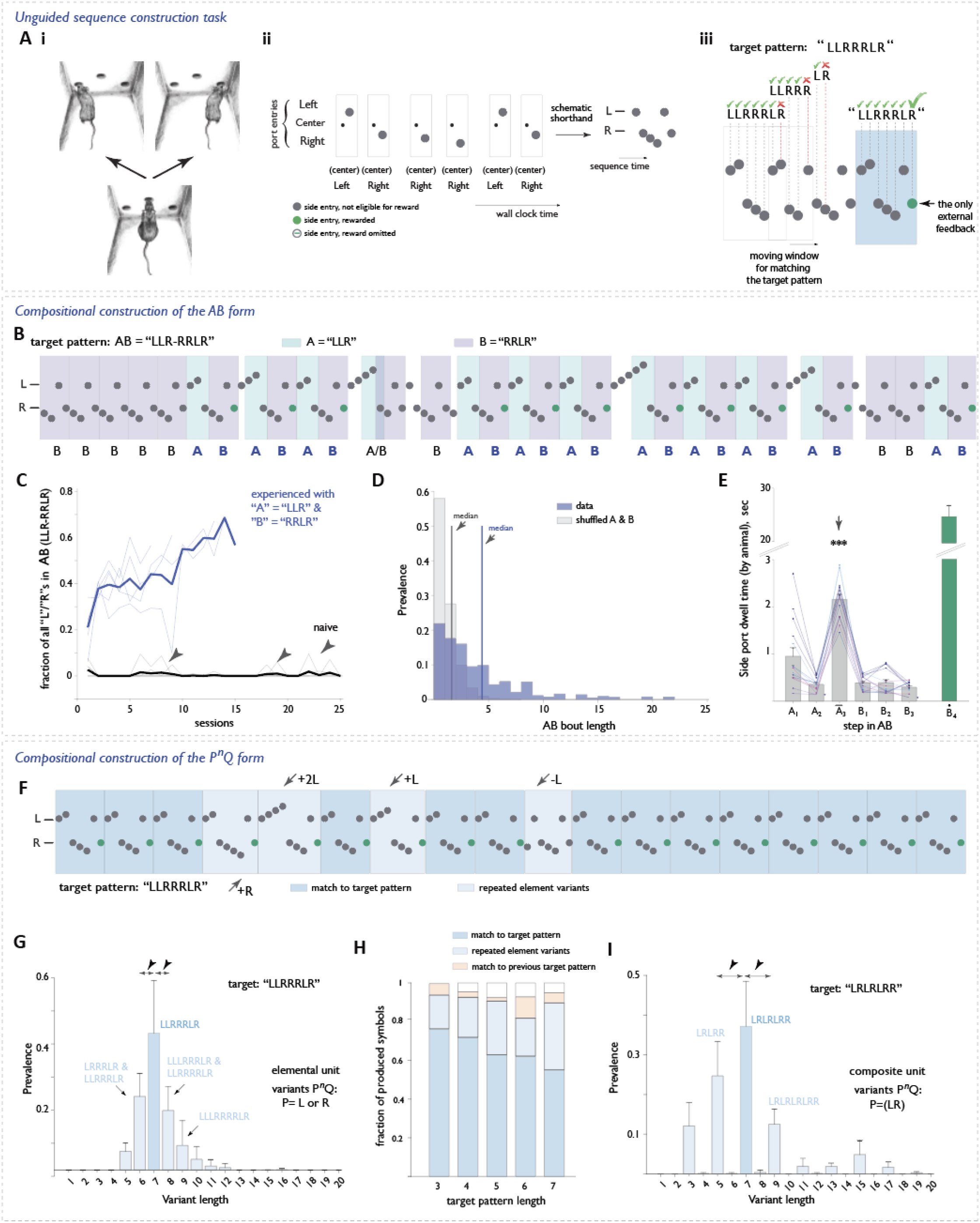
Rats innately utilize compositional construction in an unguided sequence discovery task. **A**, Task concept. **i**, Animals are placed in an impoverished environment with left, center, and right nose ports. **ii**. Only two elemental action units can affect the state of the world: L, defined as a center port poke followed by a left port poke, and R, defined analogously with a right poke. The only feedback is reward delivery when an unsignalled, experimenter-defined target pattern (here, “LLLR”) is detected in the recent behavioral history (at the side port on the last step). From the animal’s perspective, nothing indicates the composition or even the length of the target pattern, and rewards are sparse and irregular until the target pattern is discovered, and behavior is consolidated into meaningful compositions of L and R elements in the appropriate order. **iii**. illustration of a sliding window applied to the stream of behavioral actions in search of a match to target pattern. **B-E**, *Second-order compositional construction of the AB form in unguided sequence discovery*. **B**, Example behavioral trace during “LLRRRLR” learning, illustrating the A=LLR and B=RRLR components. **C**, AB learning curve, plotted as the fraction of all L and R steps. Animals with prior experience of the first-order constituents (LLR and RRLR) rapidly acquire the second-order composit. In contrast, naïve animals fail to learn despite serendipitouslyencountering reinforced matches to the target pattern (arrowheads). The stark difference between the two groups of animals in the ability to discover these longer target patterns highlights the power of hierarchical compositional expansion of one’s behavioral repertoire. **D**, Prevalence of AB bouts of different lengths in the data vs after A and B matches were independently shuffled. n=15 sessions, N=5 animals. Two-sample Kolmogorov-Smirnov test: D=0.50, n=2266 and 151766 (data and shuffle), p=0. Vertical lines indicate the median. **E**, Side port dwell times on each of the 7 steps in AB. Different colors represent different animals. The longer dwell time on step 3 marks the terminal step in A, exposingbehavioral reuse of first-order constituent parts. n=15 sessions, N=5 animals. One-way ANOVA: F=59.2, p-4.3×10^−26^, d.f.=5. Post-hoc Tukey HSD tests with Bonferroni correction; A_3_ vs A_1_: mean differenced=1.2,95% Cl[0.8,1.6], p=7.9×10^−13^ A_3_ vs A_2_: mean differenced= 1.8,95% Cl[1.4,2.2], pd.S×10^−21^ A_3_vs A_4_: mean differenced.= 1.8, 95% Cl[1.4,2.2], p=4.4×10^−21^; A_3_ vs A_5_: mean differenced.= 1.8, 95% 0[1.4,2.2], p=5.0×10^−21^; A_3_ vs A_6_: mean dlfferenced.= 1.9,95% 0[1.5,23], p=2.1×10^−22^. Data are meants.e.m. **F-l**, *Compositional construction of the P*^*n*^*Q form* **F**, Example behavioral traces showing repeat-element sequence variants (lighter blue) amidst predominant matches to target pattern (darker blue). **G**, Prevalence of repeat-element sequence variants of different lengths in sessions with the 7-step target pattern “LLRRRLR”. For this pattern, the form is L^3^R^3^LR, and thus sequences of the form L^n^R^m^LR where n ≠ 3 and/or m ≠ 2 constitute repeat element variants, indeed, animals varied their number independently, resulting In a set of variants differing by 1 step (arrowheads, cf. panel (I)). n=9 sessions, N=3 animals. **H**, Overall prevalence of repeat-element sequence varaiants for target patterns of different lengths Target sequences used: 3 step: “LLR”, “RRL” 4 step: “LLLR”, “RRBL” 5 step: “RRRLR” 7 step: “LLRRRLR”, “LRLRLRR” n = 15,14,11,9, and 18 sessions; N = 5,5,4, 3, and 6 animals for target lengths 3-7, respectively. **I**, Prevalence of repeat-element sequence variants of different lengths in session with the 7 step target pattern “‘LRLRLRR”. Sequence instances selected a) started with an L that followed an RR, and b) ended with an RR.Unlike in (G), the form of the target pattern is (LR)^n^ form of this target pattern. If compositional construction indeed uses a P^n^Q template, then the variants should be of the (LR) ^∧^nRform where n ≠3, and thus differ in length by 2 steps (i.e 3,5,9,11, etc arrowheads). If, however, the number of Ls could have been varied independently, then variants like LLRLRLRR or LLRLRR (length 8 or 6) would also have been observed. n=9 sessions, N=3 animals.

## Results

### Rats innately utilize compositional construction in an unguided sequence discovery task

To gain intuition for why the unguided sequence construction task is challenging and how a compositional mechanism makes finding solutions feasible, consider the analogy of your trying to reliably break into your colleague’s safe that has a variable-length combination lock. You see the buttons on the keypad, but you have no idea what sequence to generate, whether you are off by a little or a lot, and when to try pulling the door open. A brute-force approach becomes precipitously less effective for longer combinations. However, if you suspect that your colleague may have used some combination of shorter, meaningful patterns such as birthdays, your task becomes much easier. Here, we ask whether rats similarly exploit compositional solutions in a conceptually similar task with unknown target combinations of two “button presses” (elemental action units “L” and “R”, see Figure 1A).

Having previously demonstrated that rats readily discover short (length 3 and 4) target combinations (or patterns) comprising L’s and R’s^12^ (also, Figure S1A), we asked whether rats can reuse these learned sequences in further composition to match longer targets that are difficult to discover from scratch (in the sense of representing a tiny fraction of all possible combinations of L’s and R’s up to the target’s length). As an alternative to the brute-force search, we conceptualized *hierarchical* compositional construction in the rat behavior as utilizing “templates” comprising placeholder component parts <u>A, B</u>, and so forth that can be substituted with any units (elemental or composite) in the animal’s repertoire to produce a concrete behavioral instance. For example, the 7-step target pattern “LLRRRLR” can be efficiently constructed if <u>A</u> were substituted with LLR and <u>B</u> with RRLR, in striking contrast to a brute-force search through a set of 254 possible sequences of L’s and R’s up to length 7. Indeed, rats with prior experience of matching A=“LLR” and B=“RRLR” targets readily discovered AB=“LLRRRLR”, in stark contrast to their inexperienced counterparts that did not produce LLRRRLR sequences at any appreciable frequency (Figure 1B,C; as notation, we use double quotation (e.g. “LLR”) when referring to experimentally-defined target patterns, but omit quotes (e.g. LLR) when referring to sequences actually produced by animals). Furthermore, the prevalence of AB instances in the behavioral stream produced by the experienced group significantly exceeded chance juxtaposition of the previously learned A and B component parts, demonstrating that the 7-step sequence is a new behavioral unit (Figure 1D). To ask whether this AB behavioral sequence retained a signature of constituent parts, we turned to our previous finding that rats pause at the side port on the final (a.k.a. terminal) step of short sequences even without reward (^12^, see also Figure S1). We thus looked for such extended dwell times at the side port as a boundary mark for composite units in the behavioral stream, which reflects some internally generated “chunking” (cf.^5,16–18^) of action sequences regardless of whether in service of reward-expectation-type computations and/or intrinsic to the mechanism of behavior generation. Rats consistently paused after the third step across the LLRRRLR sequence instances, marking the end of the A=LLR constituent part (Figure 1E). Notably, the pause at the end of A when it was a part of AB was significantly shorter than when A was produced as a standalone unit (e.g. during unrewarded AA repetitions, Figure S2), suggesting a distinction between internal (at the end of each component part) versus external (at the end of the whole sequence) boundaries. Combined, these findings argue that rats constructed LLRRRLR in a compositional manner by concatenating LLR and RRLR component parts into a new whole.

In addition to the abovementioned construction via concatenation, expert rats often produced sequences that deviated from the target pattern in systematic ways (Figure 1F,G): increasing the number of repeated elements (“expansions”, e.g. LLR-**R**-RRLR; where bold letter indicates a repeated element), or decreasing this number (“contractions”, e.g. LLR(R)RLR or L(L)RRRLR; where parentheses indicate an omitted element). Such structured variations were present for a variety of target patterns (Figure 1H, see also Methods). The prevalence and the specificity of these variations lead us to hypothesize that they reflect a distinct compositional construction of the form <u>P</u>^n^<u>Q</u>, where <u>P</u>^n^ denotes n consecutive repetitions of unit P. A specific test of this “repeat variation” template corresponding to a general construction mechanism is that it should apply also to the case where P is a composite unit. We therefore trained a separate cohort of rats on the length-7 target sequence “LRLRLRR”, which corresponds to a construction of the form <u>P</u>^n^<u>Q</u> with P=LR, n=3, and Q=R. Indeed, expert sessions for this cohort comprised predominantly repeat variations (Figure 1I), suggesting that the construction process produced variations in repetition of the LR composite element. The specificity of these variants thus indicates a distinct compositional mechanism for expanding the behavioral repertoire.

In summary, we established that rats predominantly exhibit two qualitatively distinct forms of hierarchical compositional construction: concatenating (composite) units into a larger composite and varying the number of already-repeated elements in a composite unit. Building on the insight from our behavioral analyses, we next probed the underlying neural representation, focusing on the rat amFC (area 32d, Figure S3A). We used high-density Neuropixels probes to record simultaneously from large numbers of neurons (typically 200-500 per session) and investigated geometrical properties of the neural representation through a neural population perspective.

### Representational format of composite sequences comprises an orderly progression of L/R elements to a common endpoint

The internally driven chunking of behavioral sequences suggests that the underlying neural representation of a given sequence should be systematically structured such that subsequent computations can easily read out which action should be performed next and when to stop (to switch to a next sequence). We asked if the amFC implementation of this conforms to a particularly mechanistically simple representational scheme that is itself compositional^19,20^, utilizing independent directions in the neural space for the identity of the elemental action (L/R) vs the progression through the steps of the sequence, plus a sequence-general “stop” signal. Our task allows controlling for alternative sensorimotor interpretations by leveraging the design feature where every L and R action execution begins at the same central nose port. Therefore, unless stated otherwise, we focused on analyzing population activity in a fixed window centered on center-port action initiation, conceptualizing the coordinated activity of the neural population as a point in a high-dimensional neural state space where each dimension corresponds to the total spike count of a single neuron (Figure S3B, Methods).

#### Repeated L/R elements within composite units are represented as distinct neural activity clusters

We began by assessing whether amFC represents the step-specific identity of repeated L/R units in composites such as LLLR and RRRL. To understand the geometry of this high-dimensional neural activity, we applied principal component analysis (PCA) to identify dimensions ordered by highest to lowest variance. We found that the data is very low dimensional, exhibiting a sharp fall-off in variance captured as a function of the number of principal components (PCs) (Figure S3A), with all PCs after rank 10 compatible with gaussian noise (Figure S3D, see Methods). To denoise the high-dimensional space for analysis purposes, we therefore discard the higher, non-signal PCs (11 and beyond). Unless stated otherwise, we perform quantifications using this denoised ten-dimensional space.

Visualizing just the first three PCs revealed that amFC population activity associated with repeated executions of both LLLR and RRRL composites is organized into a set of discrete, well-separated clusters that matched the individual sequence steps (Figure 2A). We asked if the representation of a given step could be linearly separated from that of every other step. One possibility is that amFC distinguishes L versus R elemental actions^12^, but does not further separate R’s or L’s according to their position in the sequence (*Ho*, Figure 2B i, no separability of steps R1, R2, and R3 in RRRL). Alternatively, amFC could separate each step even for repetitions of the same elemental action (*H*_*1*_, Figure 2B, ii, full separation of steps R1, R2 and R3 in RRRL, we use subscripts to indicate the ordinal index in the sequence). The data much better matched the latter hypothesis, such that the step-specific unit identity could be determined from the neural activity on an action-by-action basis to an accuracy of ≥88% for all steps (Figure 2C, Methods). amFC activity is thus highly consistent across behavioral repetitions of the steps in a given composite sequence, and furthermore uniquely represents each of these steps even when they involve the same motor actions. We also found that amFC representation maintains only the steps necessary to specify a behaviorally chunked sequence, as opposed to representing all steps until/from a reward^21^ (Supplemental Note 1, Figure 4). amFC representations thus constitute both sufficient and necessary information for identifying individual sequence steps, consistent with the *self-organization* of elemental actions into composite units.

**Figure 2.**
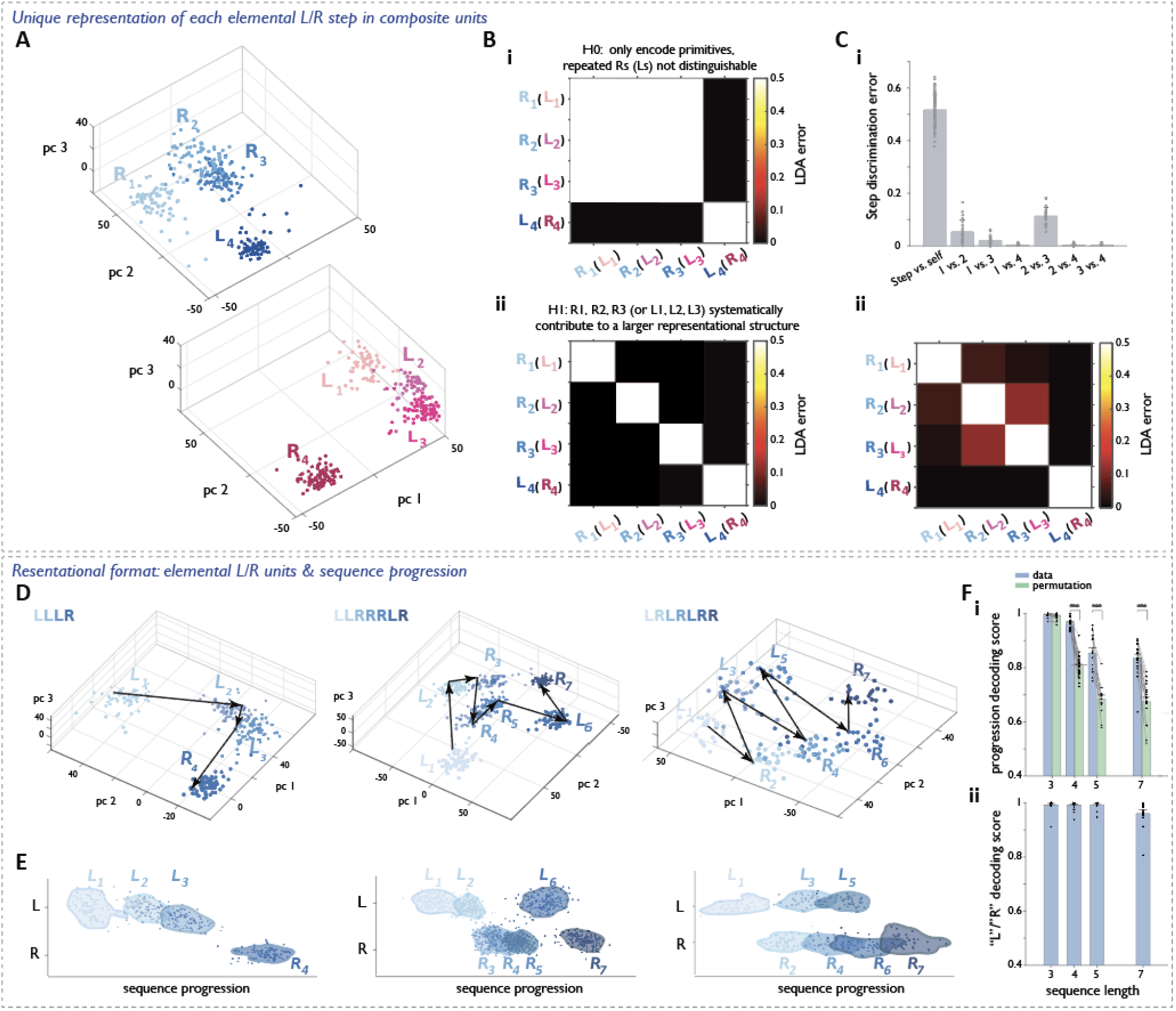
Representation of composite units in amFC permits linear readout of elemental (L/R) unit progression. ***A-C*** *amfC uniquely represents each of the composite sequence steps even when they involve the same motor actions* **A**, Population activity corresponding to constructed RRRL (top) and LLLR (bottom) composites from the same example session, visualized in the space of the top three PCs. Each point corresponds to population activity at the center port of a single step, colored by the position within the corresponding composite sequence unit. **B**, Idealized confusion matrix for the performance of a linear classifier trained to distingusih neural activity associated with different sequence steps. **i**: Performance under *H*_*o*_ assumes that only L and R can be distinguished. **ii**: Performance under *H*_*1*_ assumes that each step can be distinguished even when they involve the same motor actions **C**, Data arc more consistent with *H*_*1*_. **i**: Step-to step classifier performance. Comparison to 0.5: Wilcoxon rank sum: Step vs. self: W = 2160, p = 0.40, n = 88; 1 vs. 2; W = 0, p = 4.01 ×10^−5^, n = 22; 1 vs. 3; W = 0, p = 3 99 × 10^−5^, n = 22; 1 vs. 4; W = 0, p =1.68 × 10^−5^, n = 22; 2 vs. 3: W = 0, p = 4.01 x 10^−5^, n = 22; 2 vs. 4: W = 0, p = 1.98 x 10^−5^, n = 22; 3 vs. 4: W = 0, p = 1.68 x 10^−5^, n = 22. Data are mean +/−SEM. **ii**: Average confusion matrix **D-F** *amfC representational format permits linear readout of L/R component step progression* **D**, Example neural geometry for sequences of different lengths and composition shown from three separate example sessions, visualized in the space of the top three principal components of population activity. Activity clusters are colored by ordinal sequence step. Arrows connect neural states corresponding to successive steps within a single sequence instance, highlighting an explicit progression structure. **E**, Same data as in (d), visualized in a two-dimensional subspace defined by the direction that best captures sequence progression and the best residual L/R separation direction. The representational geometry permits a direct linear readout of both ordinal position and elemental action identity at each step of the self-organized sequence. **F**, Linear readout of L/R and sequence progression. **i**: Progression decoding accuracy for sequences of different lengths in the space of all 10 contributing Pcs. Each point compares the empirical decoding accuracy from a behavioral session with the mean accuracy obtained after all possible ordinal label permutations, ii: L vs R decoding accuracy for sequences of different lengths. See also Extended Data Fig. 5. Two-sided Wilcoxon signed-rank test: Length 3: W = 36, p = 7.81 x 10^−3^, n = 17, Length 4: W = 325, p = 1.23 x 10^−5^, n = 25; Length 5: W = 105, p = 1.22 x 10^−4^, n = 14; Length 7: W = 209, p = 1.03 x 10^−4^, n = 20. Data ore mean +/−SEM.

#### amFC representation of composite units permits <u>linear</u> readout of progression through L/R component steps

We next investigated whether the amFC neural clusters for sequences of various lengths and compositions exhibit further organization by the L vs. R identity and the sequential order/progression of steps (Figure 2D). The existence of *linear* readouts, i.e. simple weighted sums of the activity of different neurons, would point to a particularly simple mechanism for organizing the representation of a composite unit^22^. We thus searched for a two-dimensional plane in the neural state space onto which the neural activity could be projected, such that the order of steps within a sequence could be best decoded along the first axis (“progression direction”), and the residual L/R identity of each step could be best decoded along the second axis (Methods). Visualizing the high dimensional activity by projecting onto these two dimensions revealed a strikingly simple organization of neural clusters during execution of a behavioral sequence, where one can easily read out which elemental action should be produced at which step (Figure 2E). Quantitatively, this corresponds to high classification performance for both decoding the sequence step using the progression direction, and for decoding the L/R element identity using the *L*/*R* axis (Figure 2F; for clarity, we italicize neural cluster labels). These decoding performances were significantly above chance for all target sequence lengths and were not compatible with a random encoding of steps^23^, as the latter inherently has no ordering (Supplemental Note 2, Figure S5). The amFC representation thus goes beyond generic separability of steps in the sequence, and is systematically organized in a way that permits linear readout of the specific ordering and identity of L/R elements in the sequence being executed.

#### Composite units converge to a common endpoint in neural state space

Finally, we asked whether there is a signal for the end of a composite sequence that is general across different sequences, e.g. a common endpoint location in the neural state space. We visualized the representations of both L_1_L_2_L_3_R_4_ and R_1_R_2_R_3_L_4_ sequences as the rats first exited the center port and then subsequently entered the side port. At the center port, the activity patterns within the three dimensions that captured the largest variance were clearly separated for the two terminal steps (Figure 3A i, blue and orange arrows corresponding to L_4_ and R_4_, respectively). However, as the animals approached the side port, the activity patterns for the terminal steps L_4_ and R_4_ converged towards a single location (Figure 3A ii, blue and orange arrows). This convergence of activity patterns was also apparent when the data were visualized within a two-dimensional plane that spans the two sequence progression dimensions (Figure 3B). This convergence commenced as soon as the animals began to pull out of the center port on the terminal step of each sequence (Figure 3C) and thus could not be explained by a common sensorimotor experience tied to reward delivery and consumption. Rather, the observed change in the neural representation geometry suggests an internally computed reorganization to a common endpoint (e.g. related to the completion of a self-organized sequence and/or reward anticipation anticipation^16,20,24–27^).

**Fig. 3.**
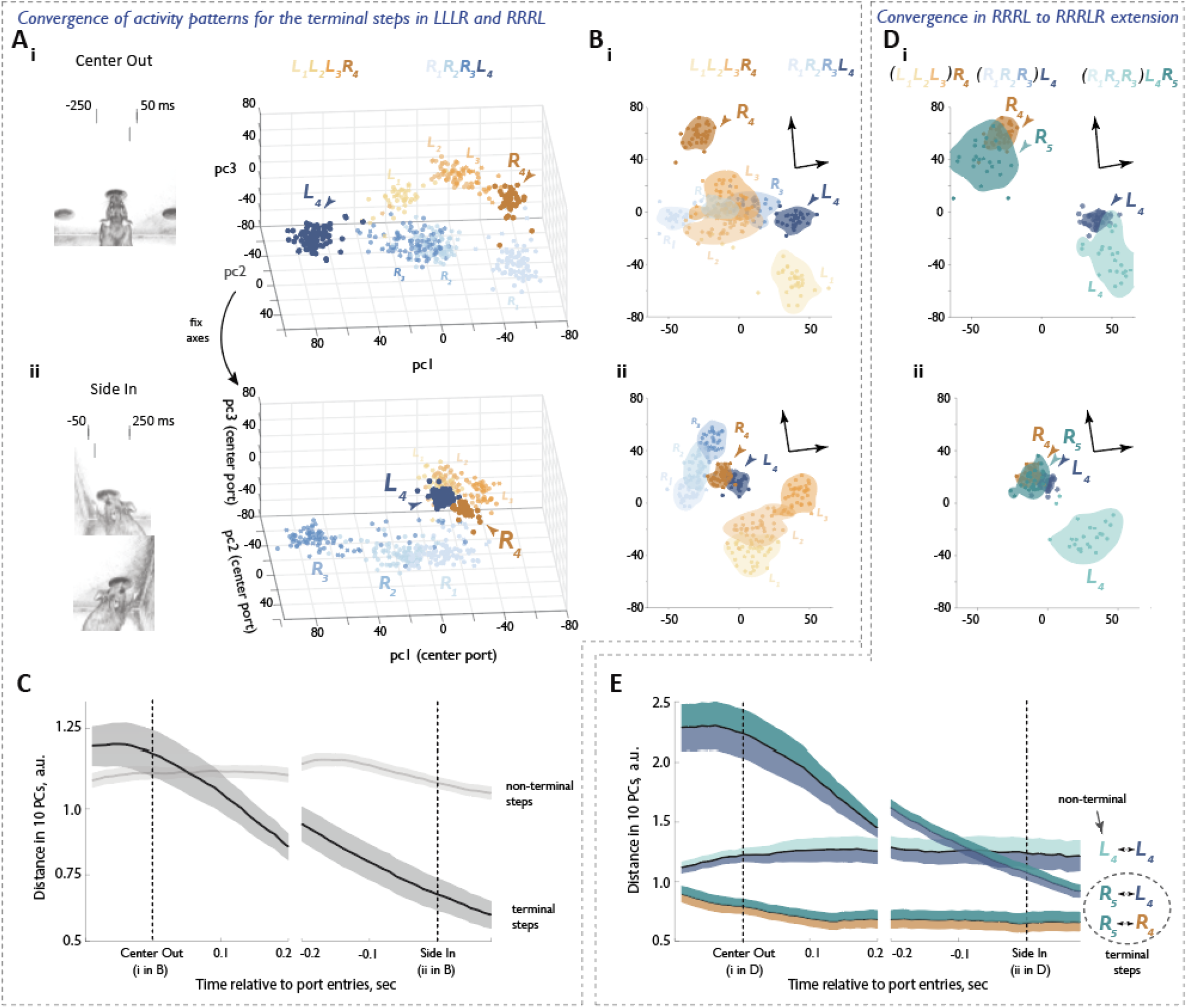
Composite units converge to a common endpoint in neural state space. **A-C**, *Convergence of activity patterns for the terminal steps in LLLR and RRRL* **A**, Example neural geometry for *R*_1_,*R*_2_,*R*_3_,*L*_4_ and *L*_1_, *L*_2_, *L*_3_,*R*_4_ during cither center port (i) or side port entry (ii). In both cases, activity is plotted in the space of the top 3 PCs of neural activity determined for the 300 msec window at the center port (i.e axes were locked before projecting side port activity; this was done to permit visualisation of the convergence for the two terminal step activity clusters (arrowheads) as the animal completes the sequence in the sideport. **B**, Neural geometry of the same representation as in (A), visualized, instead, in a plane defined by an orthonormal basis and spanning the *R*_1_*R*_2_*R*_3_*L*_4_ and the *L*_1_*L*_2_*L*_3_*R*_4_ progression directions. **C**, Change in distance in high-dimensional activity space between cluster centroids for the corresponding steps in *R*_1_*R*_2_*R*_3_L_4_ and *L*_1_*L*_2_*L*_3_*R*_4_ as animals move between the center and the side ports. The observed convergence is specific to the terminal steps indicating a common endpoint. Data are mean +/−SEM. n-15 sessions, N-5 animals. **D-E**, *Extending RRRL to RRRLR shifts the step for which activity pattern converges to the common endpoint*. **D**, Neural geometry of the terminal (*L*_1_*L*_2_*L*_3_)*R*_4_ and(*R*_1_*R*_2_*R*_3_)*L*_4_, as well as (*R*_1_*R*_2_*R*_3_*L*_4_)*R*_5_ and (*R*_1_*R*_2_*R*_3_) *L*_4_ (*R*_5_). The terminal *R*_5_ cluster, but not the non terminal *L*_4_ - that is cognate to *L*_4_ - also converges to the common endpoint. **E**, Change in distance between pairs of the relevant four clusters (*R*_4_, *L*_4_, *L*_4_, and *R*_5_) as animals move between the center and side ports. Terminal *R*_5_ converges with the other terminal clusters, *R*_4_ and *L*_4_; terminal *L*_4_ diverges from its cognate, but non-terminal *L*_4_. Data are mean +/−SEM. The SEM band is colored according to the pair of clusters being evaluated. n=15 sessions, N=5 animals.

**Figure 4.**
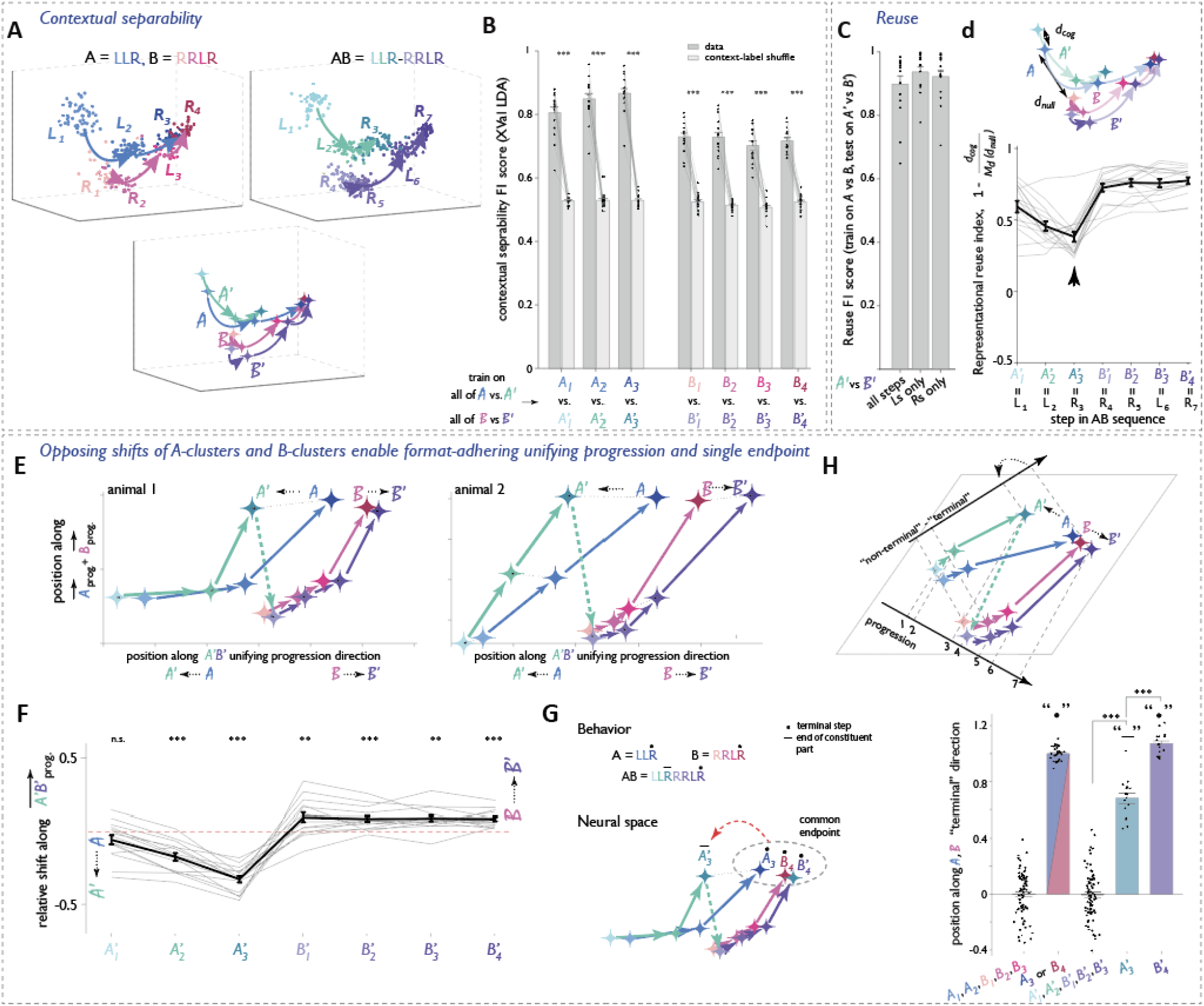
AB composites acquire a unifying progression direction via opposing shifts of the constituent part representations. **A-D** *Representations of second-order A B composites balance contextual separability and neural reuse* **A**, Neural geometry for first-order constituents (*A* and *B*) and the second-order composite ( (*A*’*B*’) in an example session, projected into the three-dimensional space that maximally separates the 14 duster centroids (visualization only). **B**, Performance of linear classifiers distinguishing the neural representations of *A* vs *A*’ and *B* vs *B*’ (in the original 10 PCspace). Two-sided Wilcoxon signed-rank tests (data vs shuffle) for each of 7 comparisons: n=15, W=120, p=6.1×10^−5^. **C**, Performance of a linear classifier trained to disnnguish *A* vs *B*, tested on *A*’ vs *B*’. Two-sided Wilcoxon signed-rank tests (data vs shuffle) for each of 3 comparisons: n=15, W=120, p=6.1×10^−5^. **D**, “Representational reuse index” (see text) for each step in AB. Arrowhead: unique displacement from *A*_3_ to *A*’_3_ (see panel (G)). **E-H** *Systematic shifts of A clusters and B-clusters enable format-adhering unifying A’B’ progression and single endpoint (see also ED Fig. 9)* **E**, Geometry of *A*’,*B, A* and *B* cluster centroids, projected onto the two-dimensional space defined by the *A’B*’ progression direction and the average of ft *A* and *B* progression directions for two animals. See also Figure S11. **F**, Displacement of individual clusters along the *A’B*’ progression direction relative to their cognate clusters in *A* or *B* across sessions.Two-sided Wilcoxon signed-rank tests (shifts vs 0); A_1_: W=36, p=0 19; A_2_: W=1, p=1.2×10^−4^; A_3_: W=0, p-6.1×10^−5^; B_1_: W=105, p=8.4×10^−3^; B_2_: W=117, p=3.1×10^4^; B_3_: W=112, p=1.5×10^−3^; B_4_ W=120, p=6.1×10^−5^ **G, Left:** Representational geometry of *A*’_3_,*B*’_4_, *A*_3_, and *B*_4_.*B*’_4_ remains near *B*_4_. and *A*_3_ within the shared endpoint zone, as expected for a terminal step; *A*’_3_ is displaced away but remains in a position consistent with its demarkating the boundary of a component part. **Right:** Normalized positions of clusters along the non terminal-terminal axis for *A* and *B* (defined such that all clusters except *A*_3_ and *B*_4_ are assigned position 0, and *A*_3_ and *B*_4_. position 1). *A*’_3_ occupies a unique position along this axis. Two-sided Wilcoxon rank-sum with Bonferroni correction: bar1 vs 3: n1=75, n2=75, W=5615, p=l, bar2 vs 5: n1=30, n2=15, W=603, p=0.15; bar4 vs 3: n1=15, n2=75, W=1243, p=4.0×10^−9^; bar4 vs 5: n1=15, n2=15, W=121, p=1.2×10^−5^ **H**, Conceptual summary of the transformation, where opposing shifts of A- and B-clusters (straight arrows) establish a unifying *A*’*B*’ progression directionand ensure a single endpoint (curved arrow).

If a common endpoint is fundamental to the organization of the neural representation of composite units, a strong prediction is that the endpoint representation should shift to the new terminal step when animals extend a composite unit to create a new whole. To test this, we trained rats experienced with the L_1_L_2_L_3_R_4_ and R_1_R_2_R_3_L_4_ sequences to also construct the R_1_R_2_R_3_L_4_R_5_ sequence and examined amFC activity patterns for all three sequences in the center port (Figure 3D i) and side port (Figure 3D ii). Consistent with the endpoint prediction, all terminal steps in all three sequences converged, including the last R (R_5_) in R_1_R_2_R_3_L_4_R_5_ (Figure 3E). Sequence representations in amFC thus seem to be constrained to converge at a common endpoint, even when modified through subsequent learning.

Could the population-level endpoint convergence be implemented by a subpopulation of goal- and reward-expectation cells^16,20,24–27^? To address this, we searched for neurons that are active at the terminal step of every sequence (Methods). Cells that were selectively active during the terminal R in LLLR and RRRLR and the terminal L in RRRL were consistently observed across sessions (Fig. S6A,B). However, these sequence-general “terminal” cells constituted only ~8% of active amFC neurons (Figure 6C). Moreover, removing these cells from the above analyses did not alter our population-level findings (Figure 6D), indicating that they do not drive the observed endpoint convergence. Instead, this subpopulation may play a distinct role such as signaling to long-range downstream targets; our findings may also point to goal/reward signaling that is implemented at the whole-population level.

All in all, the above findings establish what we refer to as the representational “format” of composite units in amFC: a linearly decodable arrangement of the elemental identity of steps in a sequence, with the last step anchored to a sequence-general endpoint. These observations argue that the mPFC population representation of composite *behavioral* units constitutes *neural* units that further compositional modifications should operate on. To resolve how format constraints are reconciled with the reuse of constituent parts in subsequent compositions, we turn to the two distinct forms of second-order compositional construction in the rat behavior.

### Geometry of second-order composition via concatenation (<u>AB</u>)

#### AB composition corresponds to contextually shifted representations that nevertheless retain high similarity to their constituent parts

When a composite behavioral unit is reused in a higher-order composition, part of its neural representation should be preserved (reflecting reuse), while a separate component should encode the specific compositional context in which it is used. To examine this for the <u>AB</u> form of composition, we selected “LLR” and “RRLR” as targets for A and B since the corresponding “AB” target pattern (“LLRRRLR”) is prohibitively difficult for animals to discover through non-compositional search (see “Rats innately utilize compositional construction in an unguided sequence discovery task”). Each session in these experiments included “A”-rewarded blocks where animals predominantly executed A sequences, from which we obtain standalone-*A* neural representations. These blocks were alternated with “AB or B”-rewarded blocks where animals predominantly produced AB sequences but also a significant frequency of standalone-B sequences outside of an AB context (Methods). As such, these experiments provided both across-block representations of A and within-block representations of B that we use as references for comparing to the neural representation of AB. Visualizing *A, B*, and *AB* in the three dimensional subspace that best separated the centroids of all neural activity clusters (Methods) revealed that the representation of the AB whole comprised two parts that appeared close to, yet distinct from, the representations of the standalone A and B references (Figure 4A). We refer to the two representational parts of the composite AB as *A’* and *B’* having verified that they are highly separable from *A* and *B* across all animals (Figure 4B; as a reminder, we italicize neural cluster labels).

We next determined how similar the representations of the original, standalone composite units A and B are to their corresponding behaviorally reused counterpart in the larger composite AB. As the simplest test or such representational reuse, we established that a linear classifier trained to distinguish *A* from *B* generalized reliably to the *Aʹ*/*Bʹ* pair (Figure 4C). Consistent with this basic indication of representational reuse, comparison of cluster-to-cluster distances in the neural state space revealed systematic similarities between the standalone and composite representations. For all steps in *Aʹ* and *Bʹ*, neural activity clusters remained closest to their respective cognate clusters in *A* and *B* (Figure S7). We quantified this representational similarity by comparing the empirical cognate distances to those expected under a null model that constructs all possible sequences from *A* and *B* clusters (Methods). The metric – “representational reuse index” – is bounded above by 1, corresponding to no contextual separation, and can fall below zero when contextual displacements exceed median distance to a randomly chosen A or B cluster (Figure 4D, Methods). The representational reuse index was strongly positive for all steps, indicating robust preservation of representational geometry (Figure 4D). The comparatively lower reuse index for step 3 was also consistent across sessions and animals, which we can explain below as related to it marking the end of the A component part (Figure 1E, Figure S2). The proximity yet non-coincidence of clusters in the composite compared to its constituent units argues that the amFC representation is compositional in the sense of maintaining a part-whole relationship beyond a simple juxtaposition of parts.

#### <u>AB</u> composites acquire a unifying progression direction via opposing shifts of the constituent part representations

How may we understand the contextual differences in AB composition as a geometrical operation by which the composite is assembled from the representation of its parts? In the simplest case, a single, randomly oriented contextual signal might simply “lift” both *A* and *B* representations into a separate dimension to create a *A*’*B*’^28^, otherwise preserving the geometry of the first-order constituents. Extending this hypothesis, there could be different, unrelated contextual signals for each of the constituents that form the second-order composite. However, such generic, randomly assigned changes might be insufficient to ensure that composites at the next level of the hierarchy maintain adherence to the proper representational format of a unit. In particular, when composing AB via concatenating first-order composites A and B, the constituent units each have their own (different) progression directions (Figure S8) and individual last steps converging to the common endpoint (Figure 3). These constraints presumably need to be reconciled for the AB composite to have a single consistent progression direction through the entirety of *A’B’*, and only the global last step remaining at the common endpoint. Indeed, the above hypotheses of unspecific random contextual signals are incompatible with the data (Supplemental Note 3, Figures S9,10), and we describe below a targeted representational transformation that takes place instead.

To understand the transformation from standalone *A* and *B* to composite *A’B’* clusters, we projected the population activity onto an alternative two-dimensional plane, with first dimension being the *A’B’* progression direction, and second dimension being an average of *A* and *B* progression directions (re-fit to be orthogonal to the *A’B’* direction, see Methods). Visualizing *A, A’, B* and *B’* cluster centroids in this plane showed that the orderly arrangement of all seven steps in *A’B’* emerged because the *A*’ clusters were systematically shifted earlier than *A* clusters, and the *B’* clusters shifted later than *B* clusters, along the composite *A’B’* progression direction (Figure 4E,F, Figure S11).

We also examined whether and how the *A’B’* composite satisfies the endpoint constraint by asking if there is evidence for a common location of last-step clusters. Using the notation *A*_i_ to indicate the neural cluster corresponding to the i^th^ step in *A* (similarly for *B*), <u>AB</u> composition entails constructing the second order composite *A’*_1_*A’*_2_*A’*_3_*B’*_1_*B’*_2_*B’*_3_*B’*_4_ from first-order composite units *A*_*1*_*A*_*2*_*A*_*3*_ and *B*_*1*_*B*_*2*_*B*_*3*_*B*_*4*_. If all sequences adhere to the same representational format, then all terminal neural clusters from all first- and second-order sequences, *A*_*3*_, *B4*, and *B’*_*4*_ — but importantly, not *A’*_*3*_ — should converge to a common endpoint. To assess this, we fit for a direction in neural state space that best separated the standalone *A* and *B* cluster centroids into “non-terminal” (*A*_*1*_,*A*_*2*_,*B*_*1*_,*B*_*2*_, and *B*_*3*_) versus “terminal” (*A*_*3*_ and *B*_*4*_) classes (Methods). We then projected the *A’* and *B’* cluster centroids onto this axis and quantified their positions, where 0 corresponds to non-terminal and 1 to terminal. As expected, the non-terminal *A’*_*1*_,*A’*_*2*_,*B’*_*1*_,*B’*_*2*_, *and B’*_*3*_ fell near 0, while the terminal *B’*_*4*_ fell near 1 (Figure 4G). Notably, the non-terminal *A’*_*3*_ occupied an intermediate position (Figure 4H), consistent with it marking an interior (part-level) as opposed to exterior (whole-level) boundary as previously predicted based on the animals’ behavior (Figure 1E). We emphasize that such intermediate positioning is *not* implied by the construction of a decoding axis using the locations of terminal vs. non-terminal steps of the standalone *A* and *B* representations, which have no interior boundaries. The geometrical positioning of the *A’B’* representation relative to the sequence-general endpoint thus closely corroborates our behavioral understanding of how animals hierarchically chunk first- and second-order compositions.

To what extent is each of the above geometrical differences required to establish the proper representational format? To answer this question, we began with the virtual *A*-*B* pairing and sequentially replaced subsets of clusters with their *A’B*’ counterparts—first substituting *A*_*3*_ with *A*_*3*_ʹ, then replacing the remaining *A* clusters with *Aʹ*, and ultimately substituting *B* with *Bʹ*—and quantified the corresponding changes in progression decoding. Decoding accuracy improved with each of these successive replacements (Figure S12), indicating that every component systematically contributes to establishing the unified *A’B’* progression direction. More broadly, our findings dispute the sufficiency of generic (a.k.a. random) contextual signals as a means by which amFC constructs representations of higher-order AB composites from their constituent parts. Instead, the observed reorganization is highly targeted, presumably to ensure that a characteristic representational format of action sequences consistently is adhered to in amFC across all levels of a compositional hierarchy (Figure 4H).

#### Representational geometry of <u>AB</u> composition is universal across animals

Given that the neural geometry involved in <u>AB</u> composition is highly nonrandom, we asked if the amFC of different animals arrived at idiosyncratic representations^29^ as may reflect inter-animal variability in behavior, unconstrained details in neural network solutions, etc. To directly compare the geometry of *A, B* and *A’B’* representations across sessions and animals, we defined two 7-vertex shapes for each of the 15 sessions: one based jointly on *A* cluster and *B* cluster centroids, and one based on *A’B’* cluster centroids. We then aligned these two sets of 15 session-level shapes from 5 animals using rigid body transformations only – rotation, translation, and mirroring, but not scaling – to minimize Euclidean distances between the corresponding ordinal positions (Figure 5A,B, Methods). As a control for the significance of alignment in high dimensional spaces, we repeated this procedure after randomly permuting step labels within each shape. Visualizing the across-animal overlay in the space of the top three principal components revealed a striking alignment of all 15 joint *A*-*B* shapes and all 15 *A’B’* shapes, but not of their permuted counterparts (Figure 5C,D, Movies 1&2). Moreover, the opposing shifts of the representational transformation from *A* and *B* to *A’B’* (see Figure 4E-H) were clearly evident in the across-animal alignment. This held true whether the *A-B* shapes were aligned jointly with the *A’B’* shapes (Figure 5E i, Movie 3) or simply visualized in the space of aligned *A’B’* shapes (Figure 5E ii, Methods). In other words, the detailed geometry of AB composites and their transformations to *A’B’* was stable not just across sessions *within* an individual animal, but also *across* animals, as if multiple animals converged onto the same solution. These observations argue for the universality of <u>AB</u> compositional construction both in the placements of neural state clusters that enact the representational format, and in the specific shifts applied by the compositional mechanism to produce the composite clusters.

**Fig. 5.**
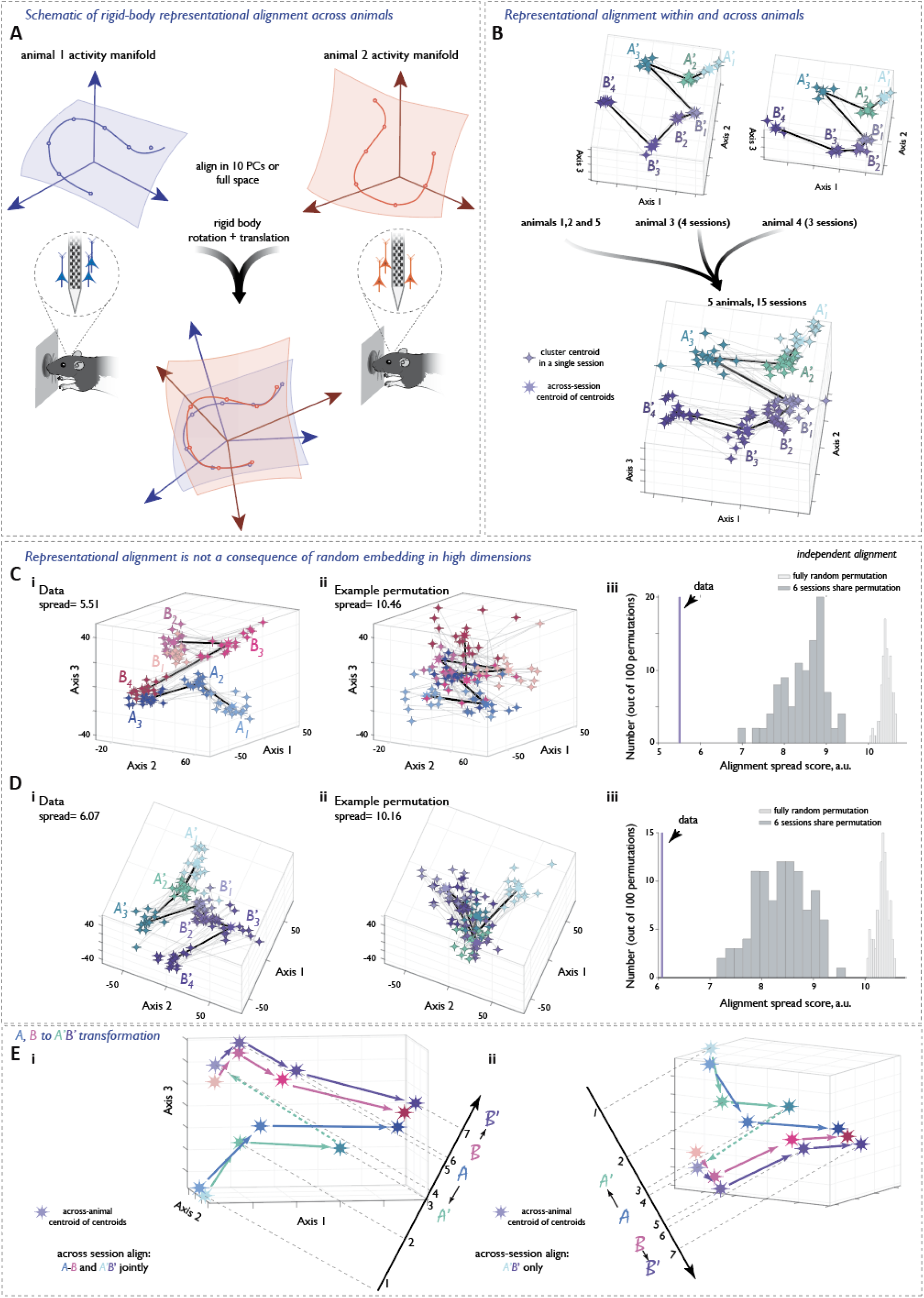
Representational geometry of AB composition is universal across animals. **A**, Schematic of representational alignment across animals. For each session, we approximate the neural manifold by defining a rigid “shape” based either jointly on .*A* cluster and *B* cluster centroids, or on *A*’*B*’ cluster centroids. The spaces are then merged across sessions and animals, and the shapes are aligned using only rotation and translation, no scaling. The alignment procedure aims to minimize Euclidean distances between the corresponding ordinal positions (see Methods). While the data in this figure are from alignments done in the space of the top 10 PCs, alignment quality is similar when done in lull (120 dimensional) space. **B**, Within-animal (top) and across animal (bottom), alignment of *A*’*B*’ shapes. Alignments were done starting from the original space of the top 10 PCs and visualized in the space of the top 3 PCs of the aligned space. See also Movies 1 and 2. **C-D** *Representational alignment is not a consequence of random emnedding in a high dimensional space* **C, i:** Global alignment of *A-B* shapes across animals and sessions. **ii**: Alignment after a single random permutation of ordinal labels within each session, **iii**: Distribution of spread (quality of alignemnt) scores across 100 fully random ordinal label permutations, and across 100 permutations where half of the sessions shared the permutation structure. The spread score observed in the data falls far outside of both distributions. p<0.01 by design (100 permutations). **D**, Same as in (C) but for *A’B’* shapes. **E**, Opposing shifts of A-related clusters and B-related clusters (Figure 4) in the relative alignment of mean *A-B* and *A’B*’ shapes. **i:** Alignment was done jointly on *A-B* and *A*’*B*’ shapes (14-vertex shapes), ii: Alignment was done on *A’B*’ shapes only. Position of each *A-B* shape in the common space was determined by applying the rigid-body transformation that was applied to the *A’B*’ shape from the same session. See also Movie 3. n=15 sessions, N= 5 animals

**Figure 6.**
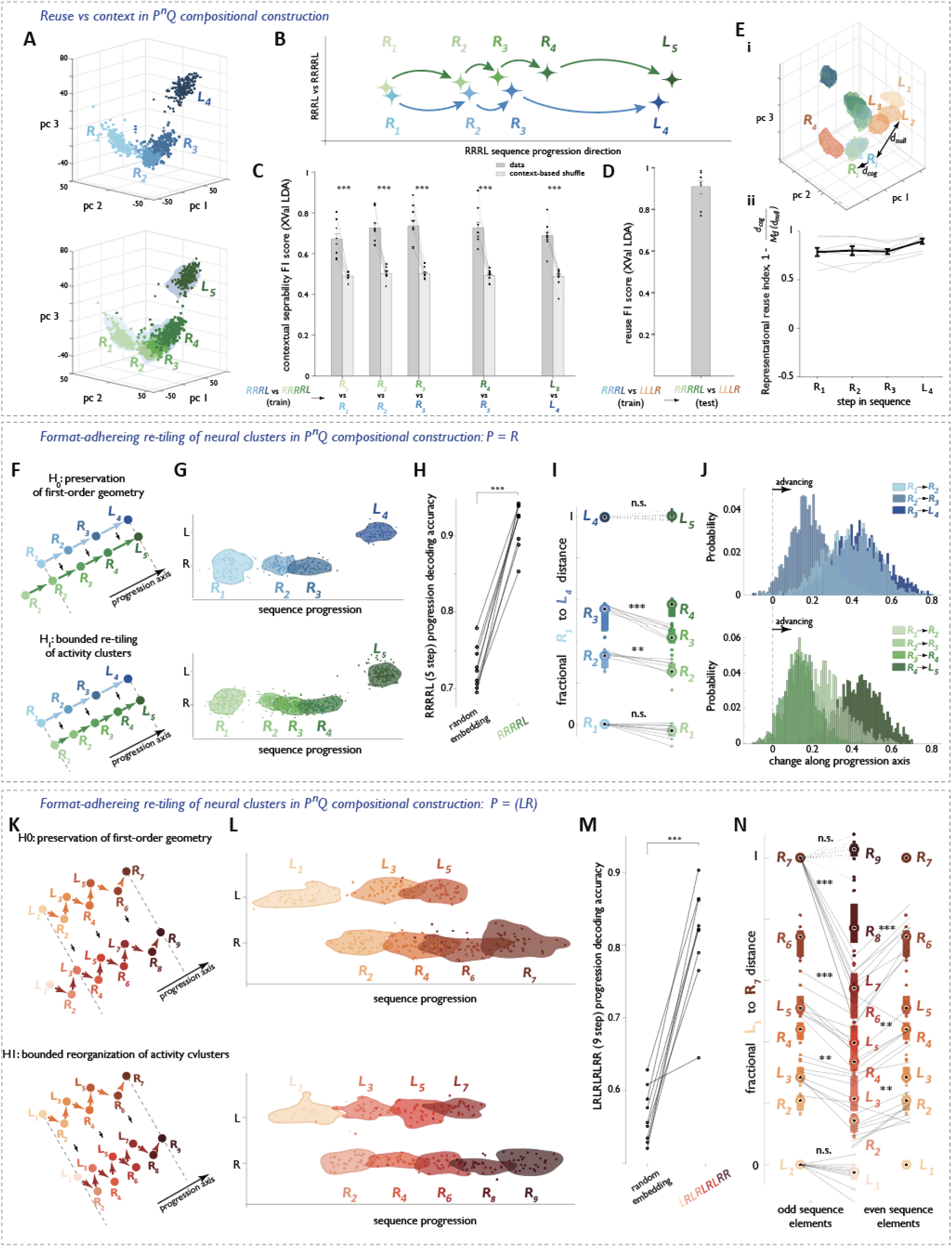
Format-adhering re-tiling of neural clusters along the progression direction in <u>P</u>^n^<u>Q</u> composition. **A-D** *Reuse vs context in P*^*n*^*Q composition* **A**, Neural geometry for RRRL (top) and its compositional expansion *RRRRL* (bottom; *RRRL* clusters overlaid on RRL cluster outlines in blue) visualized in the space of the first three principal components for an example session. Points are colored by step Identity. **B**, Same session as in (a), shown in the two-dimensional space defined by the *RRRL* progression direction and the residual direction that best separates *RRRL* and *RRRRL*. Note the consistent displacement of *RRRRL* clusters away from *RRRL*. **C**, Cross-validated Fl performance of a linear classifier distinguishing the neural representations of *RRRL* and *RRRRL*. A single classifier was trained on *RRRL* and *RRRRL*, then tested for each step individually. Two-sided Wilcoxon signed-rank tests (data vs shuffle) for each of the 5 comparisons: n=9, W=126, p =4.11×10-5. **D**, Performance of a linear classifier trained to distinguish *RRRL* and unrelated *LLLR*, tested on *RRRRL* vs *LLLR*. Robust generalization of classifier performance demonstrates representational reuse. **E**, l eft: Schematic illustrating the computation of distances to cognate clusters (d_(_cog_)_) and to clusters from unrelated sequences (d_(_null_)_, calculated using clusters for the LLLR sequence (orange) in the same session). Right: “Representational reuse index* (see text) quantifying representational similarity for each step during the transformation from *RRRL* to *RRRRL*. **F-J**, *Format-adhering re-tiling of neural clusters in P*^*n*^*Q composition (P=R)* **F**, Schematic of the predicted representational transformation from *RRRL* to that of *RRRRL* under two hypotheses. Under *H*_*0*_, expansion preserves the original geometry, yielding consistent cluster locations, and adds a clustrer outside of the original boundary along the progression direction (top). Under *H*_*1*_, expansion requires a rearrangement of cluster positions to re-establish the core representational format within the original boundary (bottom). **G**, Example session showing the core representational format of RRRL (top) and RRRRL (bottom), illustrating preservation of a shared boundary along the progression direction despite expansion. **H**, RRRRL progression decoding accuracy either expected under random embedding or observed in the data. Two-sided Wlicoxon rank-sum: W=45, p=4.11 x 10^−5^, n=9 sessions, N=3 animals. **I**, Dataset-wide normalized positions of *R*_1_,*R*_2_,*R*_3_,*L*_4_ and *R*_1_,*R*_2_,*R*_3_,*R*_4_,*L*_5_ cluster centroids projected onto the *RRRL* progression direction. For *R*_2_ vs *R*_2_ two-sided Wlicoxon rank-sum: W=54, p=1.19 x 10^-3^, n=9 sessions, N=3 animals. For /*R*_3_ vs *R*_3_. two-sided Wlicoxon rank-sum: W=45, p=4.11 x 10^−5^, n=9 sessions, N=3 animals. **J**, Step-wise displacement along the *RRRL* progression axis for all instances of RRRL (top) and RRRRL (bottom), revealing a consistent positive displacement indicative of monotonic advancement on each behavioral step. **K-N**, *Format-adhering re-tiling of neural clusters in P*^*n*^*Q composition (P=LR)* Same analyses as in (F-H), but for the transformation from LRLRLRR to its expanded variant LRLRLRLRR. For clarity, the comparison of odd step shifts and even step shifts are done separately, on the left and right side of the plot respectively.

### Geometry of second-order composition via repeat variations (<u>P</u>^n^<u>Q</u>)

Having established that second-order composition via concatenation (AB) corresponds to a targeted contextual transformation of its constituent representations, we now investigate whether the same type of format-adhering representational transformation also applies to composition via repeat variations (<u>P</u>^n^<u>Q</u>). We have already argued based on the behavioral prevalence and specificity of these variations that they arise from compositional generation and not, for example, errors or lapses in keeping track of the number of repeated elements. In addition, the highly structured nature of the amFC representations provides neural evidence of a compositional origin for this behavior (Supplemental Note 4, Figure S13).

#### <u>P</u>^n^<u>Q</u> composition corresponds to contextually shifted representations that nevertheless retain high similarity across repeat variations

We first asked if there is a distinction between the representation of RRRL and that of its expansion, RRRRL, as might reflect a part-composite relationship between the two. When projected onto a two-dimensional plane defined by the *RRRL* sequence progression direction and the residual direction that captured the most deviation away from *RRRL*, the representations of RRRL and its variant RRRRL appeared highly similar, with no apparent increase in cluster spread for the latter (Figure 6B). At the same time, the RRRRL and RRRL representations were linearly separable through a small but highly significant shift orthogonal to the progression direction (Figure 6B,C). This contextual separability is analogous to the separability of the whole vs part representations in AB composition.

We next asked whether the similarity in representational geometry of sequence variants reflects reuse. As the simplest test of such *representational* reuse, we again first verified that a linear classifier trained to distinguish the RRRL representation from that of an unrelated sequence in the same session (LLLR) generalized reliably to RRRRL (Figure 6D). To further examine the systematic similarities between RRRL and RRRRL, we asked whether the distances from the four original neural activity clusters (*R*_*1*_*R*_*2*_*R*_*3*_*L*_*4*_) to the respective closest cluster in *R*_*1*_*R*_*2*_*R*_*3*_*R*_*4*_*L*_*5*_ were smaller than expected under a session-matched null model. The null distribution was constructed by measuring distances from each original cluster to its counterpart in all possible four-step sequences randomly assembled from the LLLR clusters (Figure 6E). For each original cluster, we then quantified the “representational reuse index” that compared the distance to its closest RRRRL cluster with the median of the distances under the null model. Across steps, sessions, and animals, neural reuse indices were consistently high, and cognate-cluster distances remained low (Figure 6E), indicating systematic preservation of representational structure. Thus, this form of second-order composition is also accompanied by substantial reuse of high-dimensional neural geometry.

#### <u>P</u>^n^<u>Q</u> composites reuse progression directions via a re-tiling of neural cluster positions

The above shift orthogonal to the progression direction that separates representations of sequence variants is in principle consistent with the simple contextual “lifting” hypothesis. However, this hypothesis also predicts that distances between *RRRL* neural clusters should be preserved by operations that vary the number of repeated *R*’s. In compositional expansion, this can for example be achieved by “adding” activity clusters for repeated elements at a position before the original starting cluster (Figure 6F, *H*_*0*_). Contrary to this prediction, we observed that the positions of the neural clusters for the first and last steps in the RRRRL and RRL variants were markedly similar to those for RRRL, despite their differing lengths (see Figure 6A). Nevertheless, sequence variants adhered to the general representational format of composite units (Figure 6G), permitting robust linear progression decoding (Figure 6H), prompting us to evaluate how the positions of intervening steps were accommodated along the progression dimension.

The coincidence of the first- and last-step neural activity cluster centroids, and the consistent ordering of the intermediate-step centroids along the progression direction were highly replicable across animals and sessions (Figure 6I). In fact, the intermediate steps in the expansion *R*_*1*_*R*_*2*_*R*_*3*_*R*_*4*_*L*_*5*_ occupy a significantly expanded range within the first-last-step bounds, with *R*_*2*_ and *R*_*3*_ occurring *earlier* than their corresponding steps in RRRL, and *R*_*4*_ occurring *later* than the last *R* in RRRL (see Figure 6I, see also Figure S14 for contractions). Moreover, when we performed a timepoint-by-timepoint analysis by mapping the neural state corresponding to each action to its location along the progression direction, we found that the change in location from one step to the next is almost exclusively positive (Figure 6J), corresponding to steady advancement throughout each instance of executing RRRL variants. In other words, the stepwise ordering of neural clusters held even at the single-action level, pointing to a dynamic adjustment in how population activity progresses rather than only a statistical consistency across instantiations, and further arguing against a memory lapse origin of compositional variants. Importantly, a similar constrained reorganization characterized LRLRLRR sequence variants where expansions and contractions involved a composite subcomponent, P=LR (Figure 6K-N). These observations again argue against a simple, format-agnostic “lifting” contextual signal, and instead suggest that <u>P</u>^n^<u>Q</u> composition involves a bounded re-tiling of neural cluster positions along the progression direction, respecting the representational format of composite units. In sum, preservation of representational format across levels of compositional hierarchy is shared by both forms of compositional construction utilized by the rats.

#### Representational geometry of <u>P</u>^n^<u>Q</u> composition is universal across animals

Lastly, we asked if the representational geometry for repeat variation composition (<u>P</u>^n^<u>Q</u>) is also similar across animals. As before, we directly compared the geometry of (LR)^3^R and (LR)^4^R representations across sessions and animals by defining a 7-vertex body based on *(LR)*^*3*^*R* cluster centroids and a 9-vertex rigid body based on *(LR)*^*4*^*R* L cluster centroids, or a joint 16-vertex rigid body. As for the case of <u>AB</u> composition, the placements of neural state clusters that enact the representational format, and in the specific shifts applied by the compositional mechanism to produce the composite clusters were remarkably consistent across animals (Figure 7, Figure S15, Movies 4-6). As such, amFC compositional geometry is universal for both forms of second-order composition.

**Figure 7.**
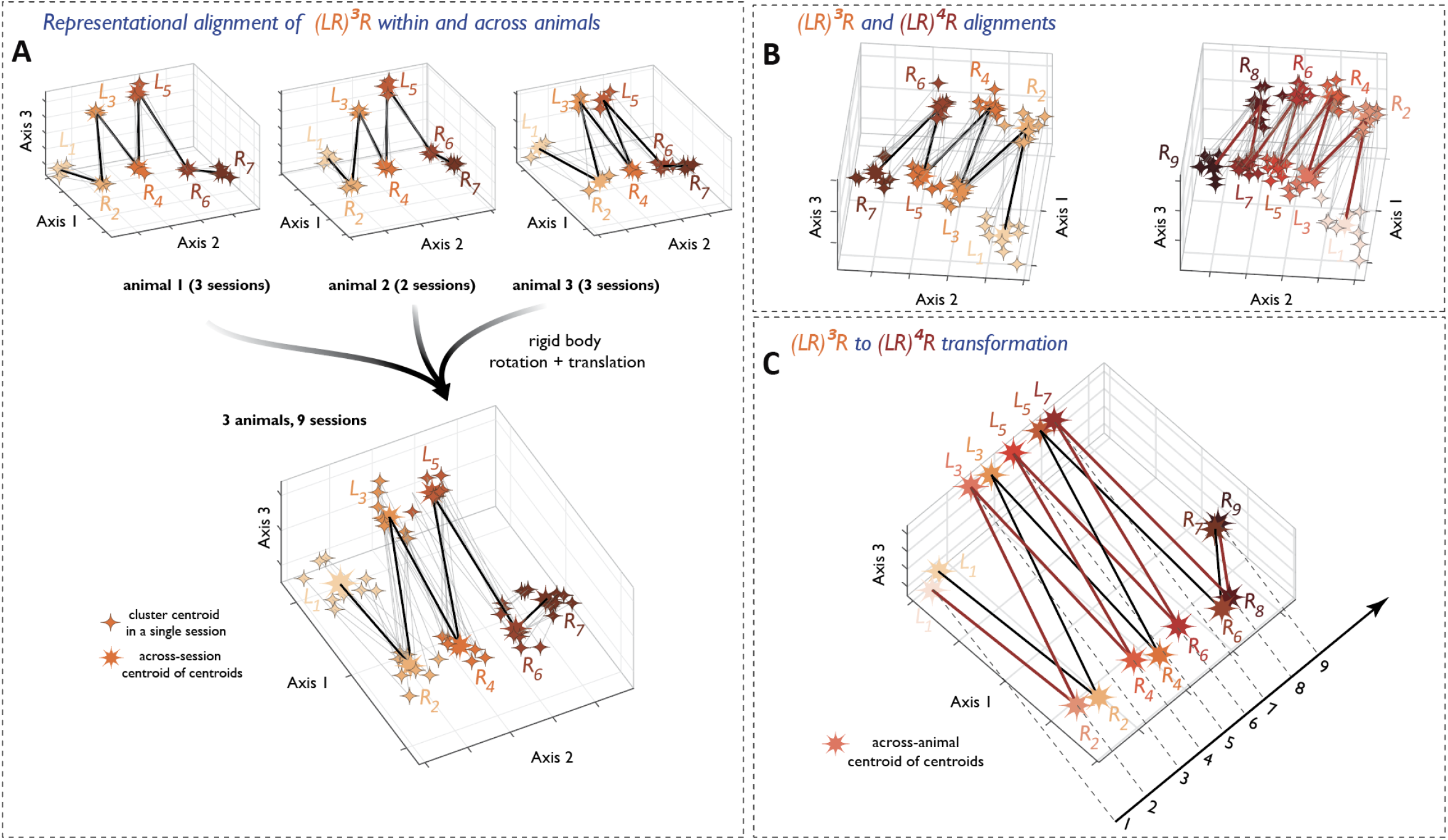
Representational geometry of P^n^Q composition is universal across animals. **A**, Within-animal (top) and across animal (bottom), alignment of (*LR*)^3^*R* shapes. Alignments were done starting from the original space of the top 10 PCs and visualized in the space of the top 3 PCs of the aligned space. **B**, Global alignment of (*LR*)^3^*R* (left) and (*LR*)^3^*R* (right) shapes across animals and sessions. See also Movies 4 and 5. C, Geometry of *LR*)^3^*R* (left) and (*LR*)^4^*R* revealed in joint alignment (as 16-vertex shapes) across animals and sessions. See also Movie 6. n=9 sessions, N= 3 animals

## Discussion

Recent years have seen encouraging progress in elucidating the neural organization of knowledge that underpins behavior, but it remains unclear how the brain systematically builds upon that knowledge to achieve an astonishing sophistication of thought and behavior across animal species. Here we identify a neural basis for hierarchical compositional construction of action sequences that enables rats to find difficult-to-discover solutions matching target sequences of unknown length and content. amFC representations of all sequences constructed by the animals adhere to a modular, low-dimensional format that encodes the orderly progression of elemental units comprising the sequence with the last step anchored to a sequence-general endpoint. A targeted representational transformation permits higher-level composites to maintain a systematic relationship to their lower-level constituent parts while providing context separation and satisfying format constraints. The identical geometry of this implementation across animals suggests what may be a universal mechanism for generating progressively more sophisticated behavioral parts to build upon prior knowledge in a changing world.

Despite longstanding interest across scientific^2,30–34^ and engineering^6,35–40^ disciplines, “composition” remains a broad and difficult to precisely define concept, and consequently, difficult to empirically measure^41,42^. Arguably the most basic assessment of whether a given system is compositional is to look for a flexible reuse, i.e. mix-and-match of parts/modules in new combinations presumably arising from an underlying compositional mechanism. Together with cellular-resolution neural population recordings and theoretical advances, this approach has in recent years made inroads into elucidating how the brain may flexibly recombine task representation subspaces^8,9^, neural state trajectories^10^, canonical object representations^43^, and dynamical motifs^11^ to produce different behaviors. In this work, we focused on the rich *construction process* afforded by compositional systems, extending beyond (first-order) signatures of reuse to seek evidence for how a system might *continue expanding* its toolkit of parts to use in subsequent compositions. Such iterative expansion requires the output of a compositional operation to be usable as an input, which in turn implies that all input and output parts should be distinct entities conforming to the same structural format required by the compositional mechanism. Indeed, in the classic example of language as a compositional system, our ability to parse and generate sentences on the fly — without the need to re-learn in a content-specific way — relies on their constituent clauses adhering to grammar rules (the “format”). We demonstrate that in compositionally generated behavior, amFC representations likewise adhere to a characteristic format across levels of compositional hierarchy, implemented through a specific and precise geometrical rearrangement of the first-order constituent parts during second-order compositional construction. Our work thus goes beyond the question of modularity to show how the amFC compositional system can make “infinite use” via iteratively combining “finite means”

Our findings highlight the remarkable relational expressivity of amFC, here shown to encode the low-level action the animal is taking, which composite unit that action is a part of, how much progress within that composite sequence has been made, and part-whole relationships in second-order composition. Recent studies have suggested that frontal areas represent remembered sequences using neural codes that are themselves compositional in nature, which may, in part, underly this expressivity ^19,20^. Our observations display some similarities with prior findings regarding the representation of first-order composite sequences, while revealing additional unanticipated structure when considering hierarchical *behavioral* compositions in open ended settings. One recent study proposed that sequence memory is implemented via compositional modules that comprise cells firing at fixed (sub-)goal/step-progression lags from an anchoring step^20^. In our data, many more neurons were tuned to progress along the *overall* sequence of goals/steps (Figure S6). One possible explanation is that the proportion of overall goal-progress vs sub-goal/state tuned neurons^20^ might be adjusted when the lack of external step-by-step feedback places greater burden on the neural system to keep internal track of self-organized sequence progress, especially during construction of more complex second-order composites. A separate study suggested that the compositional modules for sequence memory are (near-) orthogonal subspaces in the neural state space, one per ordinal rank in the sequence, although the separation between items within a rank-subspace decreased with increasing rank^19^. Intriguingly, the observed decrease in within-rank separability is compatible with our observations that all sequences converge to terminate at a common endpoint (Figure 3C). However, our observation that the neural activity clusters for individual steps in a sequence are organized along a one-dimensional progression direction implies structure beyond orthogonality in the relative arrangement of rank-subspaces, albeit longer sequences would be required for the sequence memory task^19^ to assess significance (Figure S5B). Moreover, the reuse of representational geometry in *A’B’* composites also argues against a model that invokes additional disentangled rank-subspaces for longer sequences: neural clusters in *B’* are closest to those in the corresponding first order composite *B* despite their vastly different ranks (4 to 7 vs 1 to 4). Extending the existing models to account for the constraints on the possible neural mechanisms imposed by hierarchical compositional construction would be an exciting avenue for future studies.

Future models of sequence construction will also need to account for the strikingly convergent representational geometry across animals (Fig. 5). Our observation that the neural representations of first and second-order composites can be aligned across days and animals that holds true at the geometrical level even though the numbers and identities of neurons differ for each subject extends recent accounts of representational consistency^44–46^. In our case, aligning neural data across animals only requires rotation and translation — an unexpected and unprecedented degree of similarity in the neural activity across multiple animals. The observed similarity is particularly surprising given that animals display inevitable idiosyncrasies in their learning trajectories and differences in composite sequence execution once proficient (e.g. Figure 1E). One possible reason for this universality is that the computation performed by amFC is complex and constrained to the point where it permits only a single solution^47^. More plausibly, the compositional problem has multiple possible solutions, but some genetic pre-specification of initial conditions biases the circuit to universal solutions across animals, e.g. via local or multi-regional connectivity, specific learning rules, or biophysical parameters. Investigating whether the source of the observed universality is purely computational or a combination of genetic specification and computational complexity represents an important direction for future basic research. In addition, should the universality of solutions extend to other forms of compositional thought, it could significantly lower the barrier for applying translational approaches like brain-computer interfaces in more complex cognitive settings by minimizing the necessary adjustments to decoding algorithms across subjects.

The robust compositional behavior we describe may itself have been specifically enabled by the open-ended nature of the unguided sequence discovery paradigm. Indeed, the failure of rats lacking prior experience with the first-order A and B constituents to discover AB cannot be attributed to a lack of reinforcement (Figure 1C, arrowheads) but rather points to the difficulty of credit assignment without good inductive biases^6,31^. Although the experimental logic—matching target sequences within the last seven steps—is straightforward, rats face a monumental challenge: identifying which actions in their continuous behavioral stream merit consideration^48^. Working backward from reward requires evaluating a combinatorially vast space of candidate sequences of varying lengths, a difficulty well-recognized in reinforcement learning and AI in general^49^. Biological agents routinely face such a chicken-or-egg problem: learning effective representations of the world requires constraining the space of potential causes through appropriate inductive biases, yet developing such useful biases depends on already having adequate representations^50–54^. In open-ended, unguided settings where there is little else to go on, animals may resolve this by preferentially employing compositional construction to restrict the hypothesis space by reusing previously learned relevant elements^3^. Our finding that amFC activity provides a highly structured, timepoint-by-timepoint account of compositional behavior (Figure 6J), supports longstanding neuropsychological observations of impaired ability to compose multi-step solutions to open-ended problems in amFC lesion patients^55–57^ and an emerging view that amFC plays a central role in leveraging existing structured knowledge to guide behavior in novel contexts^20,58^.

We note that the two major forms of composition we observed in the unguided sequence discovery framework may offer clues to the evolution of compositional inductive biases^59^. Composition via concatenation has a straightforward interpretation of trying previously useful behaviors in succession. Composition via repeat variation may reflect the prevalence of apparently stochastic mechanisms in nature, where the probability of an agent’s actions succeeding depends on the number of repeated attempts. Alternatively, it may represent an active search — analogous to seeking near-miss examples in one-shot learning^60^ — for the target pattern boundary. Both forms seem particularly relevant when seeking to understand the world through action^61^. Agentic learning that focuses on aspects of the world that can be affected by one’s actions simultaneously constrains the otherwise limitless possible representations and may naturally lead to a propensity to test compositions of established solutions while building an adequate representation of that world. Complementary experimental efforts that probe frontal cortical dynamics and the capacity for compositional construction in frameworks that afford the subject less agency will be informative about whether learning for control offers a useful normative explanation for observed forms of compositional inductive biases.

## Methods

### General methods

#### Subjects

All experiments were done using male Long Evans rats (400-500g). Animals were food restricted and kept at 85% of their body weight before food restriction commenced. Animals were maintained on a 12hr light/12hr dark schedule. Experiments were conducted according to National Institutes of Health guidelines for animal research and were approved by the Institutional Animal Care and Use Committee at HHMI’s Janelia Research Campus.

This study contained two main groups of animals: one group (6 animals) learned to construct LLLR, RRRL, RRRLR and their variants. 5 of these 6 animals proceeded to AB composition experiments, learning to construct the AB composite LLRRRLR and its variants (one animal dropped out because of advanced age). A second group (3 animals) learned to construct LRLRLRR and its variants. No animals were excluded from analyses: All animals (9) were included in the electrophysiological analyses.

#### Behavioral apparatus

All behavior was confined to a box with 23 cm high plastic walls and stainless-steel floors. The floor of the box was 25cm by 34 cm, and the custom-made nose ports s (https://karpova-lab.github.io/nosepoke) were all arranged on one of the 25 cm walls. All hardware was controlled and monitored with a custom-programmed microcontroller running MicroPython (micropython.org), which in turn communicated via USB to a PC running a PyControl GUI^62^ (https://github.com/pyControl/code). Nose port entries were detected with an infrared beam-break detector (IR LED and photodiode pair) or with a Time-of-Flight sensor (https://github.com/Karpova-Lab/haptic-tof-nosepoke/).. Successful port interaction was typically associated with one or more types of feedback: a change in the state of a port-embedded LED, a short auditory tone or port vibration. This was done to minimize the number of intended but unregistered sequence steps arising from port interactions that were too shallow or too fast. Note, however, that our ground truthing experiments established that omitting any feedback had little impact on the animals’ ability to discover latent structure.

Reward for eligible behavioral symbols was delivered directly at the side ports. Reward for food-restricted animals was in the form of a small volume (typically, 0.1-0.25 ml) of 10% sucrose solution mixed with black cherry Kool-Aid. Reward was delivered with the help of a custom made syringe pump (https://karpova-lab.github.io/syringepump/latest/), and dispensed for eligible behavioral steps at about 300-500 msec following side port entry.

#### Behavioral paradigm

To assess how rats intrinsically construct long action sequences and test for signatures of compositionality, we extended our previously-designed behavioral paradigm for unsupervised sequence discovery^12^. In this setting, animals are placed into an impoverished environment where only two elemental action units can affect the state of the world. Specifically, rats interact with three nose ports (left, center, and right; see Figure 1A), and the record of these interactions is used to extract two elemental action units: an ‘L’, comprising a center nose-poke followed immediately by a left nose-poke, and an ‘R’, comprising a center-poke followed by a right-poke. In this impoverished setting, rats learn to discover that different sequences of these lowest-level ‘L’/’R’ action units can lead to rewards. The specific rewarded sequence (e.g. “LLR” or “LLRRRLR”) is predefined by the experimenter, and may change randomly (in length or composition) in a block-wise fashion. Whenever a match to this prespecified sequence is detected in the behavioral record, a liquid reward is delivered at the side port entry on the last step in the sequence (for a subset of target patterns – those with the same start and end “letter”; in this dataset, “RRLR” and “RRRLR” – the sliding frame was explicitly reset upon encountering a match). Other than common sensory cues that indicate to the rat that it has successfully triggered a noseport, this sequence-specific reward is the only feedback that rats receive in the course of the experiment. From the perspective of an animal, reward occurrences are irregular and also very sparse, until it manages to discover and reliably repeat the rewarded sequence. Crucially, the requirement for center port entry to initiate every elemental action ensures that identical steps (e.g., two ‘R’s) are produced with relatively consistent movement patterns rather than varying based on prior position. This setup prevents confounding motor strategies for sequence generation and ensures that any systematic differences in neural representations of repeated Ls and Rs reflect an internal organization within composite sequence units.

Most sessions contained several unsignalled changes in the identity of the target pattern; block transitions happened ~ every 250-500 self-paced L/R actions. Given the prevalence of repeat variants, more complex target patterns often were defined in the software using regular expressions to ensure that the animals were not penalized for pattern expansions (contractions were not rewarded). For instance, for the target pattern LLRRRLR, the target pattern was usually defined as “LL+RRR+LR”, where “R+” means one or more Rs (and similarly for “L+”. Typically, a single target pattern was rewarded. One key exception to that rule was the “AB” context, which usually involved a “nested” reward contingency, with both “B” and “AB” eligible for reward. This was done both to facilitate the discovery of “AB” and to provide a within-block control for the contextual shift from B to B’ associated with second-order composition.

#### Electrophysiological recordings

A total of 9 animals were chronically implanted with Neuropixel 1.0 probes for collection of neural activity after the initial proficiency with latent sequence construction was attained. For the probe implantation surgery, trained animals were initially anaesthetized with 5% isoflurane gas (1.0 L/min) and mounted in a stereotaxic frame (Kopf Instruments). After 10-15 minutes, isoflurane was reduced to 1.5-2.0% and the flow rate to 0.7 L/min. A local anesthetic (Bupivacaine) was injected under the skin 10 minutes before making an incision. Small stainless steel bone screws and dental cement were used to secure the implant to the skull. Small stainless steel ground screw was placed above the cerebellum and connected to a wire leading to the system ground. A unilateral craniotomy (1.0 by 2.0 mm) was drilled in the skull above the site of recording and centered 3.5 mm anterior and 0.6 mm lateral to Bregma (right or left hemisphere) when recordings were targeted to the aMFC. A single Neuropixels 1.0 probe was lowered to a depth of 6.0–6.3 mm from the brain surface. The probe was permanently fixed to the skull with C&B Metabond® Quick Adhesive Cement System (Parkell) and a protective enclosure attached to skull around the probe. For the Microdrive implant, before the animal woke up, all tetrodes were advanced into the brain ~1.20 mm deep from the brain surface. Animals were allowed to recover from surgery for 10-14 days, over which time tetrodes in the one animal implanted with a Microdrive array were slowly lowered, moving approximately 40 μm/day on average. After about 10 days, animals were re-acclimated to the behavioral paradigm. When motivation and proficiency of familiar target pattern discovery regained pre-surgical levels, recording sessions began.

Data from all the animals were collected using the wireless headstage and datalogger (Neuropixels Datalogger headstage, SpikeGadgets, https://spikegadgets.com/products/neuropixels-datalogger-headstage/) powered by small Li-Polymer batteries affixed directly to the headstage. Battery rundown constrained the duration of recording sessions (typically, 3-5 hrs). 384 out of 960 channels were preselected to cover the depth of 0 to 4 mm from the brain surface and often included units in the more ventral part of mPFC (not included in this study). The identity of channels specifically within area 32d was verified post-hoc through histological analysis.

Animals self-paced their behavior and sometime took breaks (typically 5-30 minutes). Electrophysiological data were collected for a stream of 1000-5000 behavioral actions (L/R).

#### Session selection

To ensure adequate sample size for electrophysiological analyses, we selected sessions with the longest recording time and the highest prevalence of target pattern matches. The aim of maintaining the same group of animals throughout the study’s trajectory (i.e. through initially establishing stable performance on a variety of first-order sequences, following with the acquisition and stable performance of second-order composites, following sessions with both first-order and second-order composites) made the set of suitable sessions in each category rather restricted. Furthermore, the foundational comparison of representational transformation between first-order parts and second order composites required an extra degree of selectivity when choosing samples in each category (see below). However, the restricted size of recorded session best suited for these analyses was offset by the universality of representational geometry. We thus aimed to include 3 highest quality sessions (ultimately, statistical sample size) for each animal; when more sessions were available, we chose sessions separated by days to weeks to confirm that the neural geometry remained stable for each animal (max within-animal separation of 35 days).

#### Ephys data preprocessing

Raw electrophysiological data were spike-sorted using Kilosort 4 (https://github.com/MouseLand/Kilosort), which performs automated detection and clustering of extracellular spike waveforms across recording channels. The resulting clusters were then manually curated using Phy (https://github.com/cortex-lab/phy) to refine unit isolation by inspecting waveform shape, auto correlograms, refractory period violations, and cluster stability over time. This combined automated and manual approach yielded a set of well-isolated single units suitable for longitudinal analyses of amFC population activity.

All neurons that passed manual curation after automated spike-sorting were included in all analyses. The neuron numbers per session were as follows: AB cohort - 121, 146, 261, 340, 340, 340, 147, 136, 140, 122, 231, 231, 231, 200, 200; P^^n^Q cohort-199, 191, 237, 250, 250, 240, 198, 250, 250.

#### Data Analysis

##### All analyses were performed using custom scripts written in MATLAB (MathWorks)

Distinct data streams – behavioral event registration, time-of-flight signal from nose ports, video feed and electrophysiological data – were synchronized under a single clock. Four key time stamps were established for each low-level action (L/R): time of center port entry, time of center port exit, time of side port entry, and time of side port exit. If the IR beam was broken, or time-of-flight threshold exceeded, more than one time while the animal interacted with a particular port, the first timestamp in the series was taken for entries and the last time stamp for exits. No restrictions were imposed on the port dwell time or the time between center and side port entries or on the detailed trajectory taken between the ports, thus some variation in the kinematics of action execution was present in all animals.

#### Behavioral analyses

To quantify the fraction of all low-level L/R actions within AB composite, we used the “B”/”AB” behavioral blocks and separately found all matches to A=LLR and B=RRLR. Identifying these matches separately was particularly enabling given the prevalence of repeat variants, including contractions at the junction of A and B. We then calculated n= the total number of As and Bs in an immediate apposition and divided the number of low-level L/R actions within them (~n* 7(steps)) by the total number of L/R actions in the behavioral stream.

##### For dwell time analyses, however, we focused exclusively on exact matches to LLRRRLR

Prevalence of matches to target and repeat element variants for distinct target patterns was quantified by 1) categorizing each low-level action (L or R) as belonging to an instance of a particular class, 2) computing the fraction of all actions in the contextually-appropriate part of the session that falls into each of the classes. “Match to target pattern” and “repeat element variant” were defined according to the form of the experimenter-defined latent target pattern, For instance, for “RRRL”, the form of the target pattern is R^3^L, and thus sequences of the form R^n^L with n greater (or less) than 3 are repeat-element variants (e.g. “RRRRL” an expansion and “RRL” a contraction). For “LLRRRLR”, the form is L^2^R^3^LR, and thus sequences of the form L^n^R^m^LR where n is not 3 and/or m is not 2 constitute repeat element variants. Indeed, all animals varied those repeat elements independently. Finally, animals that were trained to construct “LRLRLRR”, produce sequence variants that conformed to the (LR)^n^R form of this target pattern.

To construct the distributions of repeat element variant prevalences for the 7-step target pattern “LLRRRLR” (Figure 1G), we selected all of the instances that matched the pattern L^n^R^m^LR, where n >=1 and m>=2. We then calculated the number of instances for each given pattern length and normalized it by the total count. To construct an equivalent distribution for the target pattern “LRLRLRR” (Figure 1I), we selected all the instances that a) started with an L that followed an RR, and b) ended with another RR. We then calculated the number of instances for each given pattern length and normalized it by the total count.

#### Ephys Data analysis

Except when otherwise noted, all analyses of population activity were done for a fixed time window (usually 300-500 msec) aligned on center port exit. Center port exit rather than entry was chosen because animals displayed different center port dwell times. Using a fixed time window permitted us to avoiding time warping when comparing neural activity between individual sequence steps. When analysis was instead done for a time window aligned to side port engagement (see section on convergence at a common endpoint), the window instead was usually centered on port entry to exclude the period of reward delivery and consumption.

Once the analysis window was chosen, firing rates associated with all low-level actions (Ls and Rs) in the continuous behavioral stream were calculated for all isolated units (by dividing spike number by the window length). The pre-processed data for each session was then assembled into a two-dimensional matrix with individual isolated units as rows and their FRs for individual behavioral symbols as columns.

We found that the data is very low dimensional, exhibiting a sharp fall-off in variance captured as a function of the number of principal components (PCs) (Figure 3C). Therefore, to denoise this high-dimensional activity while preserving its geometry, we applied principal component analysis (PCA), and then evaluated beyond what rank, all remaining PCs were compatible with Gaussian noise. Specifically, we tested each PC dimension with a Kolmogorov-Smirnov test against a Gaussian distribution with the sample mean and variance (see p-values plotted as a function of the PC dimension in Figure S3D). Given that PCs after rank 10 were compatible with gaussian noise (Figure S3C), we discarded the higher, non-signal PCs (11 and beyond) unless stated otherwise. Thus, most analyses were performed using this ten-dimensional space of the contributing PCs.

Note that we always utilized the entire session (i.e the entire stream of L/R actions) to define the PCA space, even when that session contained multiple blocks with distinct target patterns. In contrast, the set of (L/R) actions for which population activity patterns were displayed in a given figure, or used in a specific analysis, varied depending on the question asked.

#### Linear discriminant analysis for separability of sequence steps

For each pair of sequence steps of interest, we took values of the top 10 PCs and used them as predictors for the linear discriminant analysis (function ‘fitcdiscr’ in Matlab). These linear classifiers were always trained using 80% of sequence instances and tested on the left-out data; the cross-validation process was repeated 10 times. The figures thus report 10-fold cross-validates classifier performance metrics. When the datasets were balanced, we reported classification accuracy. In the case of imbalanced datasets, we reported the F1 score.

#### Sequence progression directions and spanning subspaces

To find the best direction in activity space for ordinal decoding without enforcing equal spacing between individual steps, we used proportional odds logistic regression with neural firings as predictors and ordinal steps as the outcome. As a background test for the quality of this fit, we ran a genetic algorithm with the utility function of finding a dimension along which the ordinal steps are fully separable (all points across letters and sequences are in the correct order when projected onto this direction). The ordinal regression converged reliably, and the two methods converged to similar values.

To independently estimate the accuracy of sequence progression and L/R decoding for a given sequence (and to visualize a 2D plane that best captures progression and L/R discrimination), we first found the best sequence progression direction in the denoised ten-dimensional activity space, constructed using all low-level L/R actions in the session. We then calculated the null space of the progression vector, projected our data into this 9-dimensional space, and, finally, did a linear regression on this subspace to predict the binary L/R value. Importantly, since we used the entire stream of L/R actions in a given session, not just those that belong to the target sequence, we were able to decorrelate these otherwise coupled variables (sequence progression and L/R).

To visualize a 2D plane that best captures progression directions of two different sequences, we first independently found best progression direction for each sequence. These two vectors form a generally non-orthogonal basis in the multidimensional space. We thus normalized them and -performed a Gram-Schmidt orthogonalization to find an orthonormal basis lying in the same plane as the progression vectors. We further rotated this plane so that our progression vectors (black arrows in Figs. 3 and S8) lie roughly symmetrically around the diagonal of the new orthonormal basis. This way we visually did not favor one or the other progression dimension to occupy a privileged position of being vertical or horizontal.

#### Progression decoding accuracy score and permutation controls

After finding the progression dimension, a separate decoding score was calculated. First, we established N-1 ordered boundaries for N ordered classes, chosen to maximize the number of points of class *i* correctly falling between boundaries *i* − 1 and *i*. The boundaries were found by searching the full space, accelerated with a dynamic programming approach. As the final accuracy score, we reported Cohen’s linear weighted kappa.

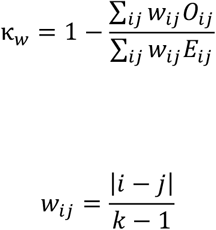

where O is the confusion matrix, E is the chance-level confusion matrix (from the marginals), and k is the number of classes.

As a control, we generated all possible permutations of ordinal positions (for example, all first steps were swapped with all second steps; N!−1 permutations total), and for each permutation found the best progression direction and the associated decoding accuracy. Decoding accuracy for the data was then compared with the average decoding accuracy of all permutations.

Note that the same classification method was also applied to evaluate the performance of L/R classifiers.

#### Random embedding controls

To establish the baseline for classification scores in the absence of sequential structure, we simulated N spherical point clusters (where N is the sequence length), distributed in a cube of n dimensions (n is the number of neurons). The dimensionality, number of points, the variance of clusters and the span of the neural space were matched to the actual data (see below).

The key parameters were estimated from the data as follows. We wanted to establish which portion of the neural space volume is occupied by clusters of activity points. To accomplish this, we first mean-centered all clusters in the data and computed the standard deviation along each dimension of the resulting pooled cluster *σ*_*cluster,i*_. Second, we computed the standard deviation of the data without centering *σ*_data,*i*_. We then computed a single ratio that pools ratios of standard deviations across all dimensions, where n is the number of neurons.

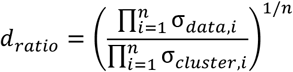

For each of N steps in the sequence, we simulated T instances (T equals to number of sequence instances in the data) across n dimensions. The center of each cluster was drawn uniformly and the within-cluster noise was isotropic Gaussian with a standard deviation s = 1/*d*_ratio_. In the end, for cluster *i*, the simulated firing rates were distributed as:

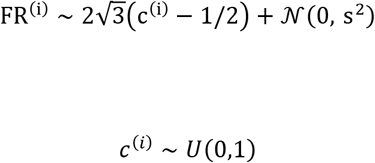

where the 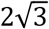 factor normalizes the uniform distribution to unit variance.

PCA was then applied to these simulated data and the regular progression fit was done. (See progression dimension fit section)For each session, the simulation was repeated 30 times and the average progression decoding score was reported.

#### Terminal cluster convergence

To demonstrate convergence of the terminal clusters as animals move from the center to the side ports, we first reconstructed the continuous neural trajectory with high temporal precision. Aligned either to the “center out” event or to the “side in” event, we binned the time with a 10 ms precision and applied a half-gaussian, strictly-causal, smoothing kernel with the standard deviation of 50 ms and the support of 500ms. This resulted in a smooth firing rate estimate for each PC, where spikes in the last 100 ms constitute 95% of the value of the estimate. Second, we calculated the mean trajectory for each step of interest in each type of sequence, and then the Euclidean distances in the 10-dimensional space between the corresponding step types. In figure 3C, distances were calculated between steps of the same ordinal value. In figure 3e, distances were calculated between specific pairs designated in the legend. Each distance was normalized by its value at the anchor point (center out, side in). Finally, we averaged the normalized distance curves across all sessions and calculated the standard error of the mean. During this step, in figure 3e, calculated distances for all non-terminal steps were pooled and averaged together without the memory of their session identity, resulting in a lower standard error of the mean due to higher N. The colors of the two SEM bands reflect the colors of the clusters, for which the distances were being evaluated.

#### Single cell analyses (Figure S6)

For each neuron, we first generated its average normalized activity “profile” within a sequence. To align its activity across different instances of the same sequence for averaging, we first calculated the median time it took the animal in that session to transition between specific events, e.g. “center-out” and “side in”. We then rounded the median duration of that time epoch to multiples of 0.2 sec. Next, for each sequence instance and time epoch, we binned that neuron’s activity into bins of 0.2sec and calculated its mean firing rate in each bin across different sequence instances. As a result, for each cell we got across-sequence-instance averaged activity that we then further normalized by the median firing rate across the time bins. We then stacked such binned, aligned and normalized activity profiles of all neurons into a matrix and resorted the rows (neurons) by the position of the bin with the maximum firing rate within each row.

We defined the terminal neurons as neurons whose mean activity for one or more sequences crossed the threshold of mean + 1.5 standard deviation only for the bins within the last sequence step. Note that a neuron was considered to have a single peak if mean activity for one or more sequences crossed the threshold of mean + 1.5 standard deviation only for the bins of a single sequence step.

#### *A’B’* neural analysis: sequence selection

Not all LLR-RRLR instances in the “RRLR”/”LLR-RRLR” context necessarily constitute second order composites: some could be the result of sporadic exploration of the “LLR” strategy within the “RRLR” context. At the purely behavioral level, these possibilities are difficult to distinguish because there are no experimenter-defined cues for “start” and “stop”, etc.

This, of course, can be evaluated based on endpoint convergence, but that would constitute a circular strategy when selecting LLR-RRRL instances for neurophysiological analyses. To avoid circularity while having high confidence about having selected second order composition instances on the basis of behavior alone, we filtered LLR-RRLR instances based on their local frequency. If the gap between two LLR-RRLR instances was less than 7 actions, the two instances are considered “nearby”. Only those instances with at least one nearby neighbor were used for analyses. Computationally, this was simplified by the 1-d “open” and “close” operations along a binary stream. With steps of LLR-RRLR instances labeled as 1, and everything else as 0, 0-runs below length 7 (small gaps) were “closed” (i.e., filled with 1), and subsequently, 1-runs below length 14 (i.e., standalone LLR-RRLR instances) were “opened” (excluded). This gave us a mask to select the with-nearby-only LLR-RRLR instances. For standalone LLR and RRLR, we just selected all rewarded LLR instances, and all RRLR instances that were not preceded by a LLR.

As we describe in the manuscript, many rats deployed repeat element expansions, including in the AB experiment (e.g. **L**LLR-RRLR or LLR-**R**RRLR). We included such variants into our analyses, leveraging our knowledge of how neural geometry accommodates repeat element expansions. For example, imagine we are trying to parse a behavioral stream “R_1_L_2_L_3_L_4_R_5_R_6_R_7_L_8_R_9_”. If we were to search with a fixed template “LLRRRLR”, we would have selected L_3_L_4_R_5_R_6_R_7_L_8_R_9_ as the steps to use for analyzing the geometry of neural activity clusters, omitting L_2_. However, from the analyses of neural geometry (see Figs. 6 and 7), we know that repeat element expansions preserve the locations of the first cluster. In this case, L_2_ defines that starting cluster in neural state space, not L_3_. Omitting L_2_ thus would introduce a substantial error into the calculation of the sequence progression direction. In contrast, L_3_ and L_4_ are squished very close together in neural space, and using just one of the two within the larger sequence is sufficient to find a good sequence progression direction. Therefore, we refined our selection method by labeling the requisite 7 steps as R-L@L@L?L?R@-R@R@R?R?L@R@, where

- “@” indicates that the preceding step is to be included in the analysis
- “?” permitted but optional repeat element expansion
- the bracketing “R” without an @ indicates that we pick the first L after an R

In other words, we selected LLR-RRLR and its repeat element expansions and matched the steps from each start of repetitions (and similarly for LLR and RRLR standalone instances).

#### Contextual separability between *A, B*, and *A’B’*

Contextual separability was assessed independently for the *A*-clusters and the *B*-clusters. In each case, we first balanced the sample size between the first-order and the second-order contexts by subsampling the larger class (subsequently repeating the random subsampling procedure 100 times). We then calculated the cross-validated LDA decoding score for each of the top ten PCs. Only those PCs yielding >50% accuracy were kept for the subsequent steps. With all selected PCs, we did one final cross-validated LDA between two classes: all clusters in *A* vs all clusters in *A’* (and similarly for *B* vs *B’*).

To determine classification accuracy for individual steps, we calculated -- for that step--the F1 score= n_correct_for_context1/ 2*n_context1 + n_correct_for_context2/2*n_context2. All of the steps were repeated for 100 different subsamples of the larger class and the average across these 1200 subsamples was reported as the final score. The same procedure was applied to a control with randomly shuffled contextual labels.

#### Representational reuse index

We used cluster centroids (median along each dimension) to calculate cluster-to-cluster distances. For a certain step i in *A’B’*, reuse index was calculated as 1 - (d_cog)i / median(d_null), where (d_cog)i is the distance from cluster i to its cognate cluster in *A* or *B*, and d_null are distances from cluster i to all clusters in the “other” first-order part, i.e in *A*, if I was between 4 and 7, and thus a part of *B’*, or to all clusters in *B* if i was 1 through 3, and thus a part of *A’*.

#### 2D visualization of *A,B* −> *A’B’* transformation

To visualize the *A,B* −> *A’B’* transformation in a 2D plane, we first determined the best *A’B’* progression vector, denoted as v_a’b’, in the space of the top ten PCs. We then removed that direction, and in the remaining 9D null space separately calculated the progression directions for *A* and for I, denoted as v_a and v_b respectively. For the 2D visualization, we used v_a’b’ as the x-axis and the vector average (v_a + v_b)/2 as the y-axis and projected cluster centroids were onto this 2D plane.

#### Calculation of opposing shifts along the A’B’ progression direction

For each pai of cognate clusters, we calculated the shift along the *A’B’* progression direction (see above) and normalized it by the distance from *A’*_*1*_ to *B’*_*4*_ along the same direction.

#### Evaluation of different models for the contextual shift in AB composition

One key difference in the way these analyses were done is that we further restricted the starting space to the top five PCs. This choice was made given that one of the key constraints on the representational transformation from *A* and *B* to *A’B’* is the establishment of a unifying progression direction through the entirety of the 7 steps of AB. In general, one can get decent linear decoding scores with random embeddings as long as the number of dimensions exceeds n-2 where n is the length of the sequences (see, for example, Figure S5C). Furthermore, we noticed that the actual shifts between *A*/*B* and *A’B’* were largely restricted to the first 5 PC dimensions. We therefore imposed this harsher restriction when evaluating to what extent simple, generic contextual shifts during second-order AB compositional construction would be compatible with the representational format constraints (Figure S9).

##### Random embedding and progression decoding scores were calculated as above

To compare the robustness of linear readout across conditions, we first matched the sample size between *A, B* and *AB* sequences by subsampling the larger groups (repeating the random subsampling 100 times and averaging the score in the end). To directly compare the outcome of simulations to the actual data, i.e., the progression decoding score of *A’B’*, we worked with the *A’B’* clusters rather than *A* clusters and *B* clusters but first shifted *A’B’* clusters in such a way that their centroid locations matched the centroid locations of their cognate *A* and *B* clusters. We denoted these clusters as *O_A*_*1*_ to *O_A*_3_, *O_B*_*1*_ to *O_B*_*4*_. We then applied simulated shifts to *O_A1* through *O_B4* in different manner for H_0_ and H_1_. Throughout simulations, each cluster was moved as a whole, i.e. without changing the within-cluster variance. After any simulated shift, we calculated the progression decoding score as above. Simulations were done independently for each session.

##### For *H*_*0*_, we simply applied the same shift vector to all seven clusters

For *H*_*1*_, we applied one shift vector to the A-related clusters, and a different vector to B-related clusters. These vectors were design to match key aspects of the data in magnitude but be otherwise randomly oriented within the 5PC space. Specifically, the starting A-shift vector was set to the average of *A*_*1*_ to *A’*_*1*_, *A*_*2*_ to *A’*_*2*_, or *A*_*3*_ to *A’*_*3*_. Its direction was then randomized by multiplying to a random rotation matrix. A similar process was applied to the B-shift vector. The first vector was applied to *O_A*_*1*_ through *O_A*_*3*_, and the second to *O_B*_*1*_ to *O_B*_*4*_, and then the corresponding progression decoding score was calculated. For each recorded session, the H_1_ simulation was repeated 100 times, giving a distribution of progression decoding scores that could be compared to one observed in that session for the actual *A’B’*. We further analyzed the relationship between the simulated shift angles and the decoding score (Figure 10).

#### Contextual separability for repeat element variants

Contextual separability between RRRL and its repeat element expansion, RRRRL, was calculated in the same manner as for A vs A and B vs B in AB composition (see above)

#### Representational reuse index for repeat element variants

Using neural activity cluster centroids for all steps in LLLR, RRRL and RRRLR, we calculated the representational reuse index as follows. For each step in RRRL, we calculated “d_cog_” as the Euclidean distance to the closest cluster in RRRRL (we took the closest, rather than the ‘cognate’ cluster in this case, because it is impossible to tell which *R* in RRRRL originated from the individual *R*s in RRRL). For d_null_, we took the median Euclidean distance to the neural activity clusters in the unrelated LLLR sequence. Finally, for each step, we calculated the representational reuse index in the same was as for AB composition, namely, as 1-d_cog_/ d_null_.

#### Retiling of cluster positions in repeat element variants

For each session and sequence set (RRRL/RRRRL or LRLRLRR/LRLRLRLRR), we first calculated the progression direction of the original sequence (RRRL or LRLRLRR). Then, for each sequence step, we projected the corresponding cluster centroid of the original sequence onto that direction and linearly warped the values such that the centroid for the first step was at zero and that for the last step at 1. We then projected centroids for all steps in repeat-element variants onto the same direction and displayed their locations relative to the [0:1] interval.

#### Step-wise displacement along the progression direction (Figure 6J)

For each session and sequence type (RRRL and RRRRL), we first calculated the progression direction (see above). Then, for each sequence instance, we projected the corresponding cluster points onto that direction and linearly warped the values such that first step of the sequence was 0 and last step at 1. For each pair of adjacent steps, we then calculated the difference (shift) along the direction. The probability distributions shown summarize these data across all sequence instances and sessions.

#### Across-animal alignment of neural geometry

Within- and across animal alignment of neural representations was done using generalized Procrustes analysis (GPA). For each session, we calculated a neural “shape” with centroids of individual steps within the relevant sequence(s) as vertices. Then we aligned these neural shapes across different sessions. GPA was only allowed to implement rigid body transformations such as translation, rotation, and mirroring, but no scaling transformations. GPA can, in principle, be performed on full neural space. However, because an N-point shape spans at most an (N−1)-dimensional subspace, performing GPA in the full neural space is unnecessarily costly. Therefore, GPA was performed in the same denoised ten-dimensional space that was used for most other analyses. Note that we separately verified that all outcomes were qualitatively the same when using the full space; for shapes with 11 points or fewer, they were numerically equivalent. The final “spread” (quality of fit) was calculated as the Frobenius distance from the mean shape, divided by the number of points. Although technically, this distance can be interpreted in terms of Hz (as the transformation comprises an unscaled rotation of the raw neural space), the scores were only used for comparison across conditions.

When multiple within-animal plots were shown side by side, GPA was used on their average shapes to align the view of all the different subplots.

Once the alignment was accomplished, a new PCA was performed on the cloud of all aligned shapes and the first 3 PCs were plotted for visualization purposes.

## Acknowledgments

We are indebted to the incredible Janelia Vivarium team for pampering the furry stars of this story. We thank Mattias Karlsson and Spikegadgets for headstage customization; Andrea Gugiu, Jon Arnold, Bruce Bowers and Bill Biddle from Janelia Experimental Technology for help with hardware prototyping, fabrication and assembly. We are grateful for particularly impactful discussions at various stages of this project with R. Axel, T. Behrens, M. Brainard, C. Brody, J. Dudman, A. Hermundstad, V. Jayaraman, K. Jensen, M. Pachitariu, S. Romani, G. Rubin and M. Sable Meyer, as well as members of the Koay and Tervo labs. This work was funded by the Howard Hughes Medical Institute.

## Contributions

MM, HW, DGRT, SAK and AYK conceptualized the study and planned out the details of the experimental design. AL carried out the engineering and software development behind the experimental ecosystem. MM, MP, HW, EK, RB and AYK performed various aspects of the experimental work. MM, MP, HW analyzed and interpreted the data with input from SD, DGRT, SAK, and AYK. SAK and AYK wrote the manuscript with input from SD and DGRT and all other authors.

## Data and code availability

All data and code will be made available upon publication.

## Supplemental Text

### Supplemental Note 1

We also investigated whether amFC representation maintains only the steps necessary to specify a behaviorally intended sequence, as opposed to representing all steps until/from a reward. To address this, we focused on instances of RRRL repetitions in the behavioral stream, where reward had been omitted for the first RRRL (Figure S4). If steps were represented relative to rewards, then amFC activity patterns for each step in the second RRRL should continue to be clearly distinguished as R5, R6, R7 and L8 respectively (Extended Data Fig. 4b). Contrary to this expectation, amFC activity patterns were closely aligned for matching steps in the two RRRL sequence instances (R1 vs. R5, R2 vs. R6, etc., Extended Data Figure 4C). Combined with the clear separability of amFC representations for the individual sequence steps, this finding shows that the internal representation of steps in amFC is consistent with the *self-organization* of elemental actions into first-order composite units, as opposed to the organization being triggered by external feedback.

### Supplemental Note 2

Is the ability to find a progression direction in the neural representation (Figure 2) surprising, or might it have arisen from the sequence steps being encoded as randomly/orthogonally separated states that inherently have no ordering? We assessed this by randomly reassigning the behavioral step labels of the neural clusters and re-fitting the progression decoder to evaluate the progression-decoding accuracy under this permutation test. To gain intuition for why the permutation test can distinguish a random arrangement from one with systematic organization, consider four clusters labeled by step (“1”, “2”, “3”, “4”), either randomly arranged on a plane or constrained to an idealized progression direction, i.e. a single line (Figure S5A). For randomly arranged clusters, permuting step labels will generally not preclude finding a direction along which projected clusters are ordered according to step labels (Figure S5A i). In contrast, it will generally be much harder to find such a readout direction after label permutations for clusters arranged along a line (Figure S5A ii). In our data, the progression decoding accuracy after permuting step labels was consistently lower for all sequence lengths (Fig. 2F i, Figure S5B), compatible with the latter hypothesis that the data consists of intrinsically low-dimensional and linearly orderable clusters. Moreover, repeating the permutation test after randomly arranging variance-matched clusters within the same space resulted instead in little to no change in progression decoding (Figure S5B; see Methods), further supporting the interpretation that the data are not compatible with a random arrangement of clusters in a high-dimensional neural state space. Also of note, the accuracy of progression decoding for the longest (7-step) sequences exhibited little decrease when restricted to the subspace formed by only the first five, or even just three, top PCs (Figure S5C). This stood in contrast to a marked sensitivity of decoding accuracy to restricting the subspace under random embedding hypothesis (Figure S5C), as expected since a linear arrangement is much lower-dimensional than a random arrangement. Combined, these findings point to a systematic organization of the amFC representation that permits linear readout of the specific ordering and identity of L/R elements in the sequence being executed.

### Supplemental Note 3

When investigating the representational transformation standalone *A* and *B* to composite *A’B’* clusters, we first assessed the hypothesis that composition corresponds to a simple contextual “lifting” of the constituent representations into a separate dimension of the neural state space (Figure S9A, *H*_*0*_). Since this hypothesis shifts every neural cluster by the same contextual vector and leaves inter-cluster distances unchanged, it is equivalent to constructing a “virtual” composite-*A’B’* dataset by where the relative geometry of clusters in *A’B’* matches that of *A* and *B* clusters (*A’B’*_*H0*_, Methods). We visualized this virtual-*A’B’* activity by re-fitting the progression and L/R directions, which revealed poor separation of sequence steps (Figure S9B i). Quantitatively, cross-validated accuracy of progression decoding through all seven steps of *A’B’*_*H0*_ (Figure S9C, *A’B’*_*H0*_) was no better than that obtained from a *random embedding* of seven variance-matched clusters (which have no intrinsic ordering, Figure S9C, “random embedding). In contrast, the *actual A’B’* composite data exhibited clear separation of individual steps along the progression direction (Figrue S9B ii), corresponding to markedly better progression decoding accuracy (Figure S9C, *A’B’*_*data*_). Thus, second-order composition entails more than a simple contextual lifting signal.

We next asked whether better *A’B’* progression decoding could be obtained by a more complex form of contextual tagging—one that shifts all A-related clusters in one common direction and, independently, all B-related clusters in another (Figure S9D, *H*_*1*_). Such a transformation would correspond to a part-specific (but not otherwise step-specific) contextual signal, e.g. “first part” for *A′* and “second part” for *B′*. To test this hypothesis, we computed, for each session, the average displacement vector from standalone-*A* to *A*’ clusters and then randomized its angle to construct virtual-*A’* clusters for *H*_*1*_ (similarly for the standalone-*B* to *B’* displacement vector, see Methods). We generated an ensemble of *A’B’*_*H1*_ datasets under this hypothesis by sampling 100 random shift angles per session and calculated the best-fit *A’B’*_*H1*_ progression decoding accuracy for each dataset (Figure S9E-F). The actual AB progression decoding accuracy of the data exceeded the *H*_*1*_ distribution’s confidence interval in one third of the sessions (Figure S9F), indicating that even a constituent-specific, but otherwise random, contextual shift was insufficient to explain the representational geometry of the *A’B’* composite in the data. Furthermore, we found a strong inverse correlation between the *A’B’*_*H1*_ progression decoding accuracy for a given pair of random shifts, and how much those random shifts pushed A-related clusters earlier and B-related clusters later along the empirical *A’B’* progression direction (Figure S10).

### Supplemental Note 4

We considered whether behavioral variations of the <u>P</u>^n^<u>Q</u> form might arise not from compositional generation but instead reflect a lapse in keeping track of the number of repeated elements. To investigate this possibility, we took advantage of the observation that in “RRRL” target pattern experiments, rats also produced significant numbers of pattern variations (Figure 1H for 4-step sequences), i.e. expansions (RRRRL) and contractions (RRL). One possibility is that a memory lapse during RRRL execution constitutes a mental “error” that could manifest as a deviation of neural activity patterns from those associated with “RRRL” sequences. However, when visualized using the three dimensions that capture the largest variance, the locations of activity clusters appeared remarkably consistent between RRRL, its expansions (RRRRL, Figure 6A), and contractions (RRL, Figure S14A). Given this orderly representation of behavioral steps, we might expect a memory lapse associated with an expansion to manifest as revisiting a step in the internal *R*_*1*_*R*_*2*_*R*_*3*_*L*_*4*_ representation (as a reminder, we italicize neural cluster labels), such that activity patterns associated with a particular behavioral step in R_1_R_2_R_3_R_4_L_5_, say R_2_, would sometimes fall in the underlying *R*_*2*_ activity cluster, but would sometimes fall in the *R*_*1*_ cluster, reflecting repetition (Figure S13B). Similarly, the activity patterns associated with the behavioral steps R_3_ and R_4_ would sometimes fall in the internal *R*_*2*_, *R*_*3*_ or even *R*_*1*_ clusters. In such cases, we would expect to see overlap in the neural activity patterns associated with distinct steps (R_2_, R_3_ and R_4)_ in RRRRL (Figure S13B). Contrary to this expectation, neural activity patterns associated with all five behavioral steps in RRRRL could be reliably distinguished using a simple linear classifier (Figure S13C). This indicates that the amFC representation for each of the RRRL variants adheres to the core representational format, with a distinct neural activity cluster for each step. The presence of distinct representations for each sequence variant indicates that the amFC represents each variant as a distinct compositional construction, rather than the possibility that these variants arise from noise in counting.

## Supplemental Figures

**Figure S1.**
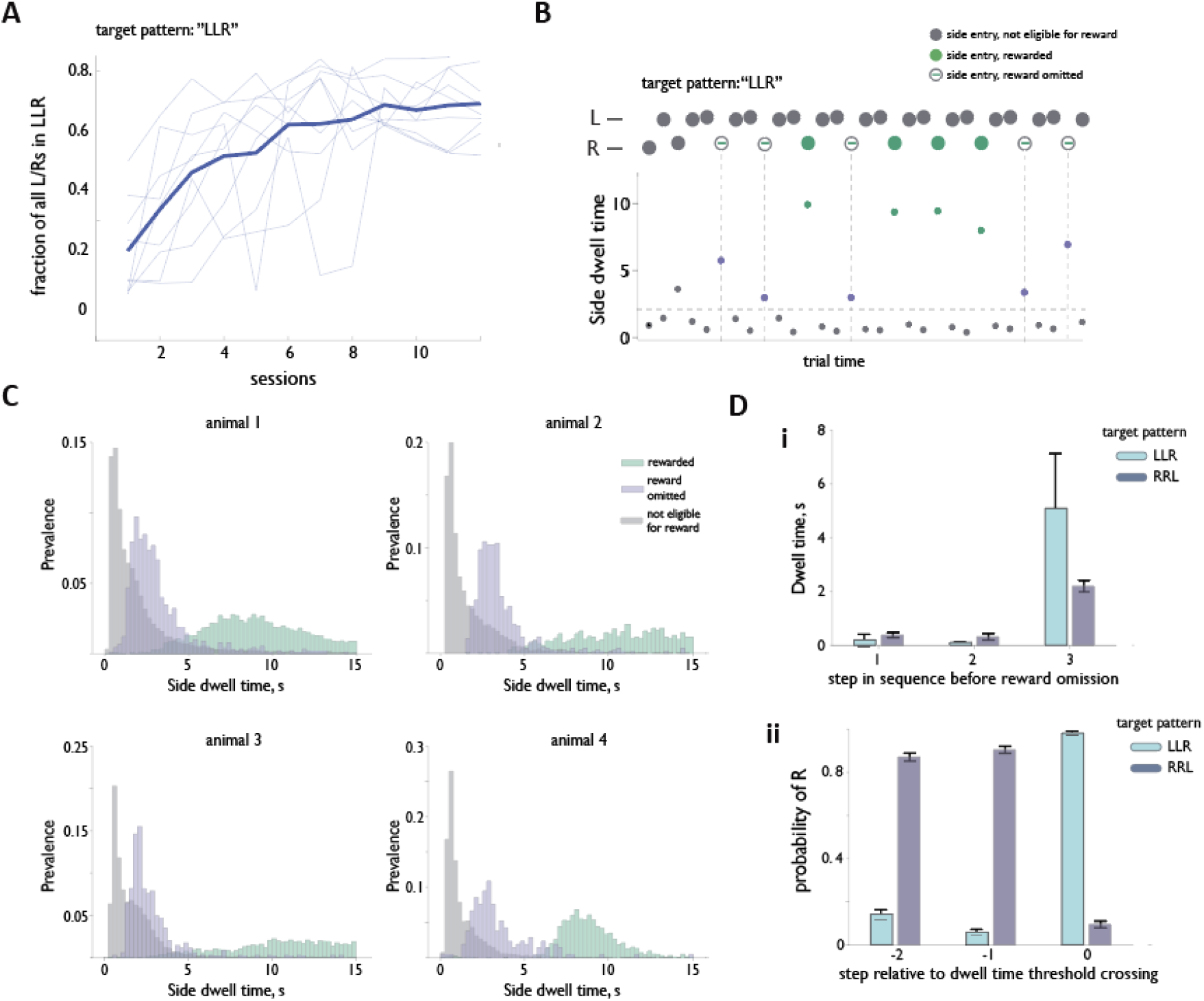
related to Figure 1 | Sequence chunking during unguided discovery of short target patterns. **A**, Learning curve for the first-order compositional target “LLR”, plotted as the fraction of all L and R steps. **B**, Side-port dwell time during an epoch in which reward is deliberately omitted for multiple correct executions of LLR (vertical dashed lines). Omissions reveal selective pauses at sequence boundaries. Horizontal dashed line indicates the dwell-time threshold used for analysis in D. **C**, Example distributions from four animals of side port dwell time for all trials in a session with alternating blocks of “LLR” and “RRL” target patterns. Dwell time wass calculated as a difference between the time of first entry into a side port and time of the last port exit within a trial. The significantly longer dwell time at the end of sequences that match the target pattern, but for which reward was intentionally omitted (purple) suggests that animals organize their elemental L/R actions into sequences. The difference between grey and purple distributions was used to find session-specific threshold for (D). Specifically, hreshold was set as 75th percentile of all dwell times for a given session. **D**, *i*, Dwell time at each step across unrewarded executions of three-step target sequences Wilcoxon rank-sum with Bonferroni correction: LLR, 1vs 2: W = 210, p = 0.08, n = 25; 1 vs 3: W = 0, p = 1.23 x 10^−5^. n = 25; 2 vs 3: W = 0, p = 1.23 x 10^−5^ n = 25. RRL, 1 vs 2: W = 379, p = 0.08, n = 33 1 vs 3: W = 17, p = 2.50 x 10^−6^, n = 3;3 2 vs 3: W = 22, p = 3.86 x 10^−6^, n - 33 *ii*, Composition of three-step behavioral histories triggered by dwell times exceeding a session specific threshold Wilcoxon rank-sum with Bonferroni correction: LLR, 1 vs 2: W = 325, p = 1.23 x 10^−5^, n = 25; 1 vs 3: W =0, p= 1.23 x 10^−5^, n = 25 2 vs 3: W = 0, p = 1.23 x 10^−5^, n = 25. RRL, 1 vs 2: W = 27, p = 5.91 x 10^−6^, n = 33; 1 vs 3: W = 561, p = 5.39 x 10^−7^, n = 33; 2 vs 3:W = 561, p = 5.39 x 10^−7^, n = 33 Data are mean +/−S.E.M

**Figure S2.**
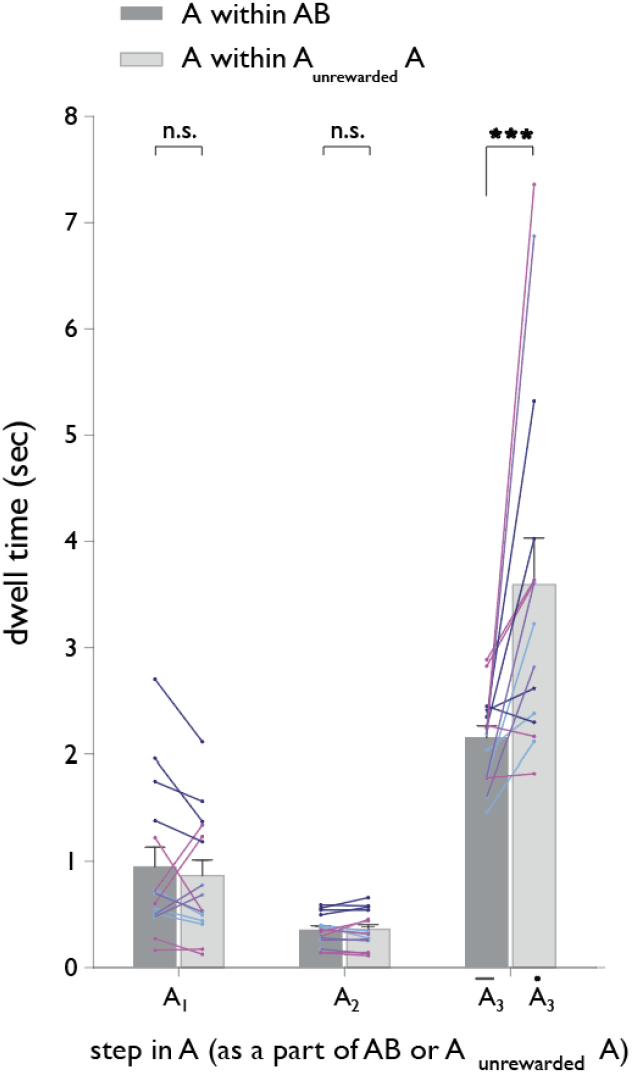
related to Figure 1 | Rats treat AB as a single behavioral unit. Dwell times for the three steps in unrewarded instances of A that are either standalone units (selected by virtue of their being followed by another A) or as a part of AA. Different colors represent different animals. n=15 sessions, N= 5 animals. Two-sided Wilcoxon signed-rank test; A1: W-statistics=74, p=0.45; A2: W-statistics=65, p=0.80; A3: W-statistics=5, p=6.10 x 10^−4^. Data are mean ± S.E.M.

**Figure S3.**
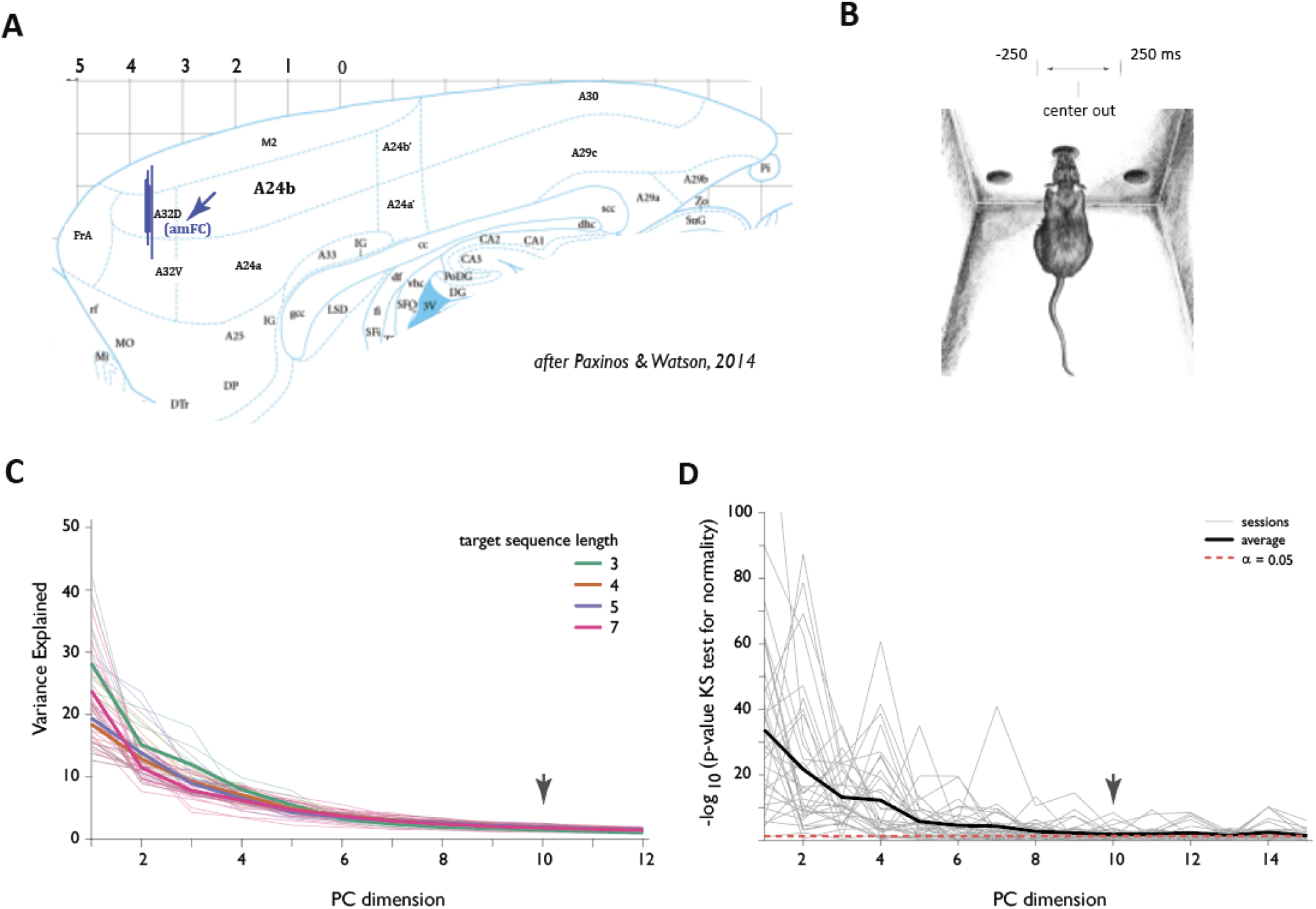
related to Methods | Details of electrophysiology experiments. **A-B**, Basics of data collection **A**, Approximate location of amFC (recording sites targeting A32D). **B**, Schematic of default analysis window alignment. **C-D**, Rationale for restricting the analyses to the top 10 principal component dirmensions of the neural activity state space **C**, Variance explained for the top 12 pricipal component directions. **D**, Normality score for the top principal component directions. PCs after rank 10 were compatible with gaussian noise.

**Figure S4.**
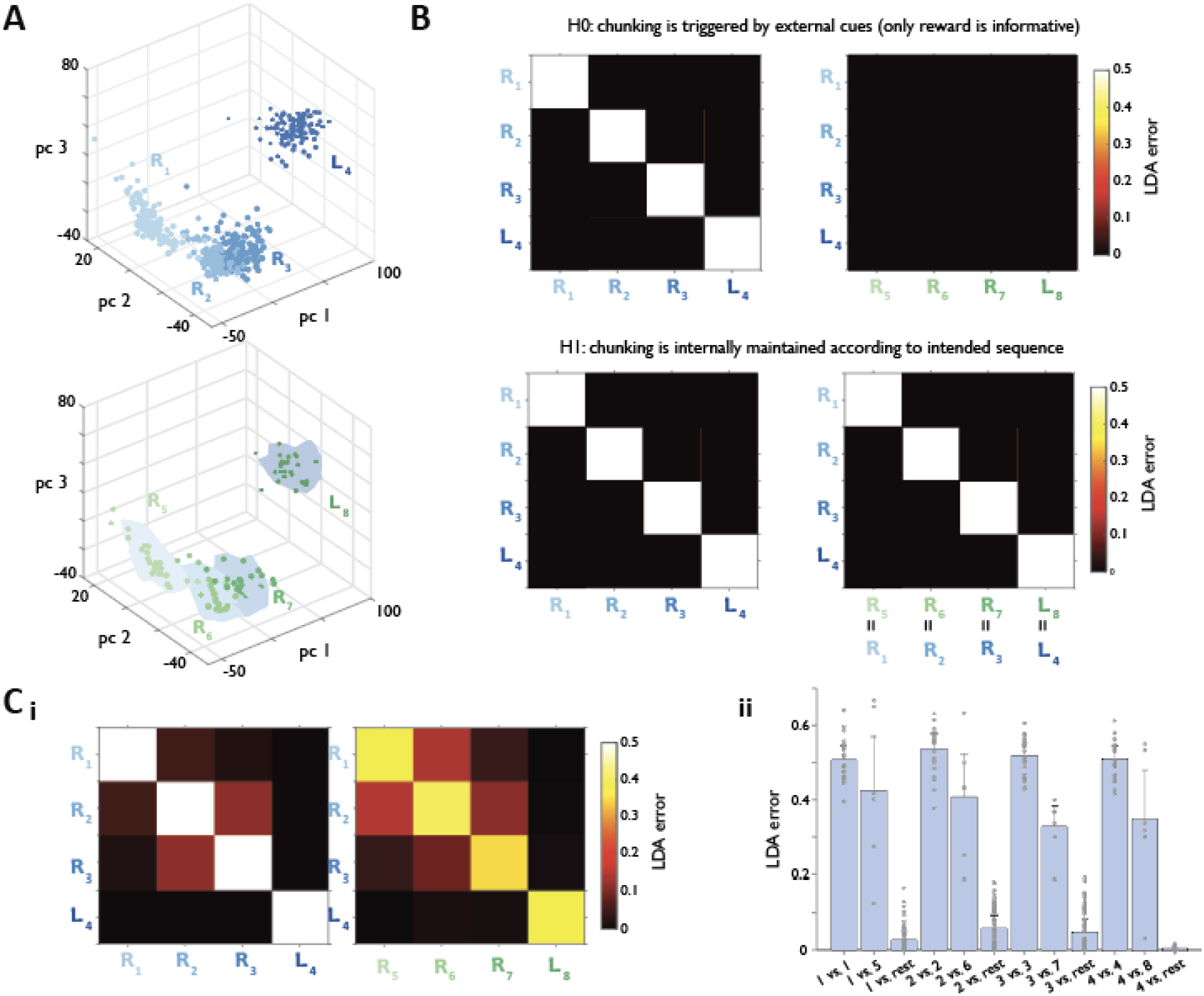
related to Figure 2 | amFC representation of composite sequences is internally organized rather than being triggered by external feedback. **A**, Neural geometry, in the same example session, of RRRL’s that follow either a rewarded (top panel) or an unrewarded (bottom panel) sequence instance. For clarity, the set of first sequence instances is given step labels 1-4, and the second 5-8. **B**, Idealized confusion matrix for vizualizing the performance of a linear classifier trained to distingusih neural activity associatedwith different sequence steps in such RRRL-RRRLs top panel: Confusion matrix under H_*0*_ that assumes that steps are represented relative to rewards; steps in the second RRRL continue to be distinguished as positions R5, R6, R7 and L8 respectively. bottom panel: Confusion matrix under H_*1*_ that assumes that steps are internally organized, making corresponding steps in the two RRRL instances indistinguishable. **C**, Data are more consistent with H_*1*_ *i*, Dataset-average confusion matrix for RRRL-RRRL instances *ii*: Step-to-step classifier performance across individual sessions. Wilcoxon rank-sum for comparison to LDA error of 0.5:1 vs. 1: W = 141, p = 0.64, n = 22; 1 vs. 5: W = 7, p = 0.56, n = 6 1 vs. rest: W = 0, p = 1.38 x 10^−12^, n = 66; 2 vs. 2: W = 194, p = 0.06, n = 22; 2 vs. 6: W = 1, p = 0.07, n = 6; 2 vs. rest: W = 0, p = 1.50 x 10^−12^ n = 66; 3 vs. 3: W = 171, p = 0.15, n = 22; 3 vs. 7: W = 0, p = 3.12 x 10^−2^, n = 6; 3 vs. rest: W = 0, p = 1.41 x 10^−12^, n = 66; 4 vs. 4: W = 145, p = 0.55, n = 22; 4 vs. 8: W = 0, p = 3.12 x 10^−2^, n = 6; 4 vs. rest: W = 0, p = 1.41 x 10^−12^, n = 66 n=6 sessions, N= 3 animals

**Figure S5.**
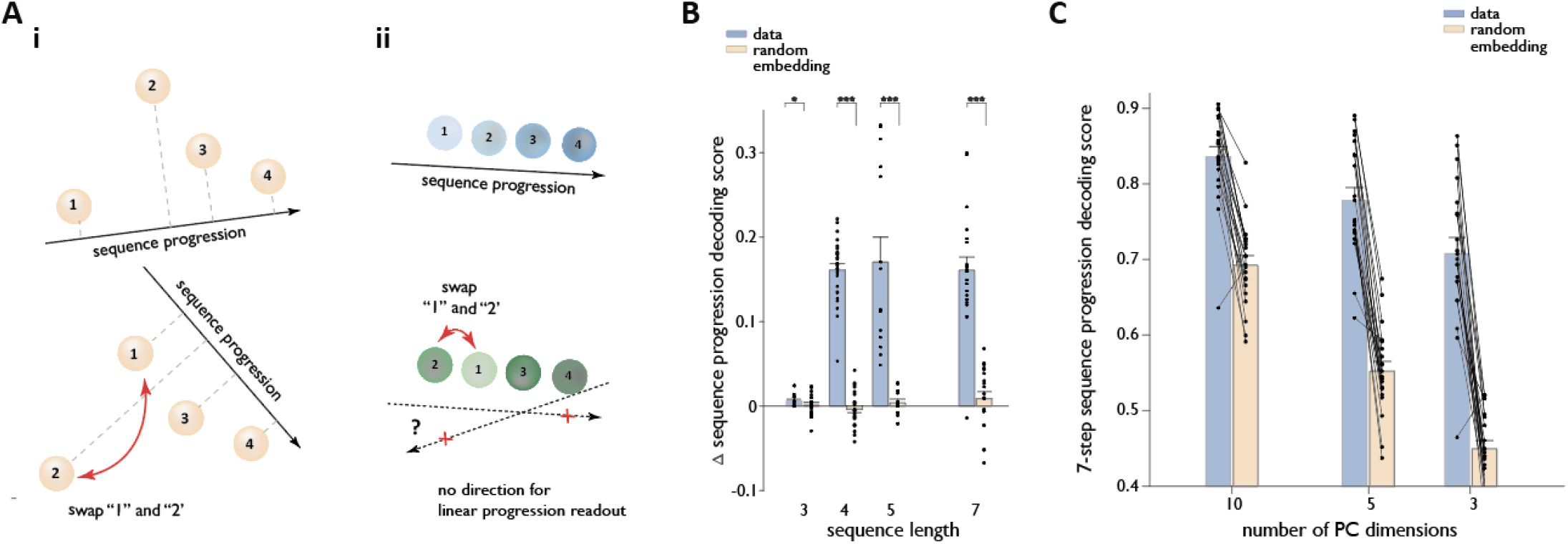
related to Figure 2 | Significance of linear sequence progression readout. **A**, Sensitivity of linear readout to ordinal label permutations in the presence of low-dimensional structure **i:** Permuting ordinal labels across randomly arranged clusters typically preserves linear decodability of sequence progression, **ii:** In contrast, assigning the ordinal label “1” to an interior cluster along the progression axis abolishes any linear readout, indicating a structured, low-dimensional organization rather than arbitrary separability. **B**, Sensitivity of decoding accuracy to ordinal label permutations for empirical data and for variance-matched random embeddings across sequence lengths. Note that sequences of length 4 or greater are needed to establish the significance of linear decoding. Two-sided signed-rank test (paired data): Length 3: W = 82, p = 0.07, n = 17; Length 4: W = 325, p = 1.23 x 10^−5^, n = 25: Length 5: W = 105, p = 1.22 x 10^−4^, n = 14; Length 7: W = 210, p = 8.86 x 10^−5^, n = 20. **C**, Decoding accuracy for 7-step sequences as a function of the number of PCs used for the dataand for a random embedding of variance-matched clusters. The modest decline in performance with dimensionality for the data is in contrast to sharp decline than for a random embedding, suggesting that linear readout is not a trivial consequence of high dimensionality. Two-sided signed-rank test: PC 10: W = 209. p = 1.03 x 10^−4^. n = 20; PC 5: W = 210, p = 8.86 x 10^−5^, n = 20; PC 3: W = 209, p = 1.03 x 10^−4^, n = 20;

**Figure S6.**
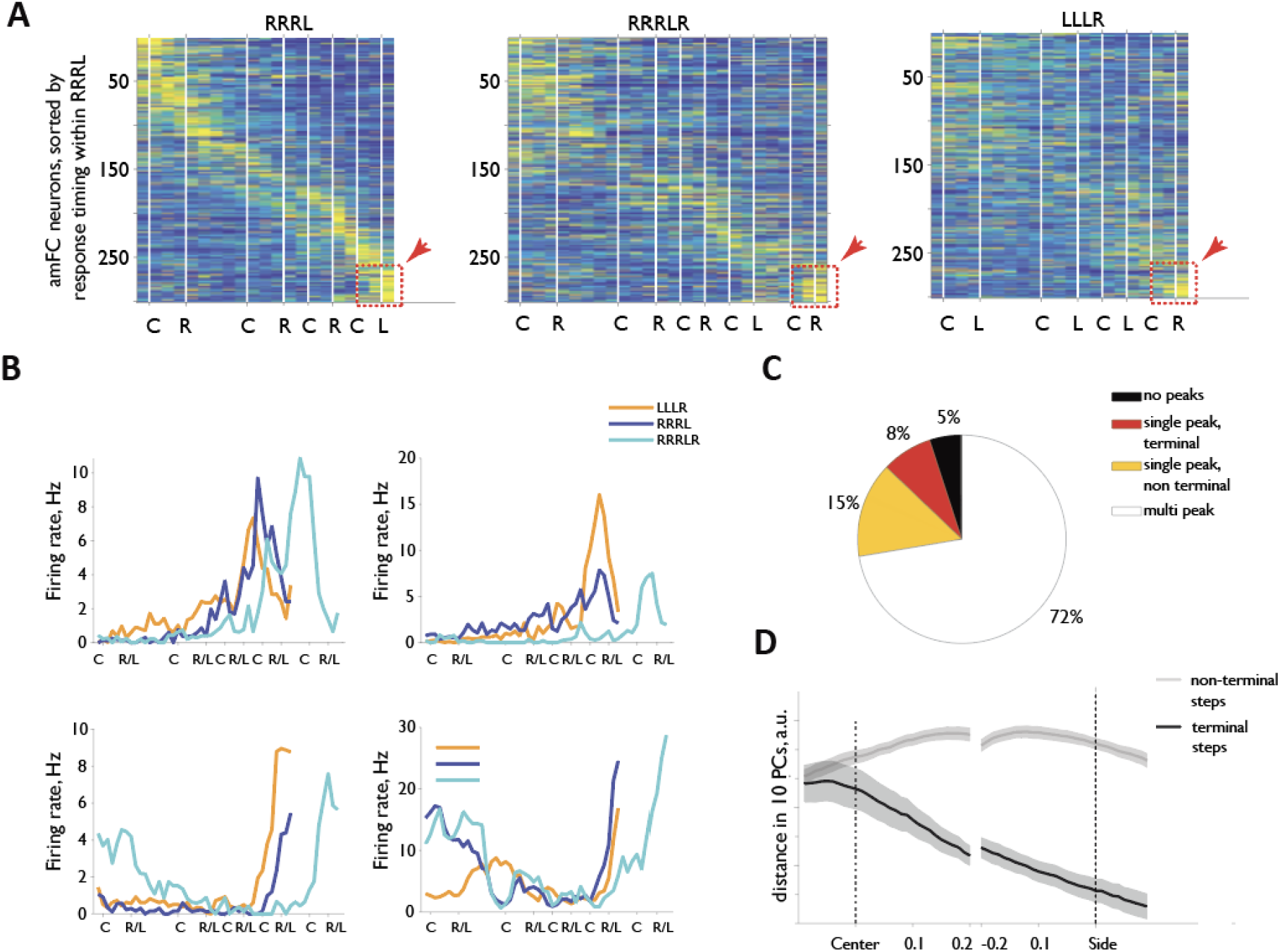
related to Figure 3 | Sequence-general ‘‘terminal” cells in amFC do not drive population-level endpoint convergence. **A**, Responses of amFC neurons during execution of RRRL RRRLR and LLLR sequences in an example session; responses of individual cells are normalized by their median (Materials and Methods). White lines represent center- and side-port entry events. Neurons are sorted by the timing of their peak firing within RRRL. Red boxes and arrowheads highlights a population of cells active during the last step of all three sequences. C: center port entry, R: rightside port entry; L: left side port entry. **B**, Average activity of four example “terminal” cells during the execution of RRRL, RRRLR and LLLR. To be included, a cell needs to show a single peak in activity in each of the three sequence that falls on the last step. In particular, the peak is on the terminal L in RRRL, but shifts to the terminal R in RRRLR. **C**, Summary of the “terminal” cell prevalence across sessions. **D**, Distance between neural cluster centroids for terminal and non-terminal sequence steps after the removal of “terminal” cells from the analysis. The unaltered convergence of the terminal clusters (cf. Figure 3) suggests that “terminal” cells do not drive this population level phenomenon.

**Figure S7.**
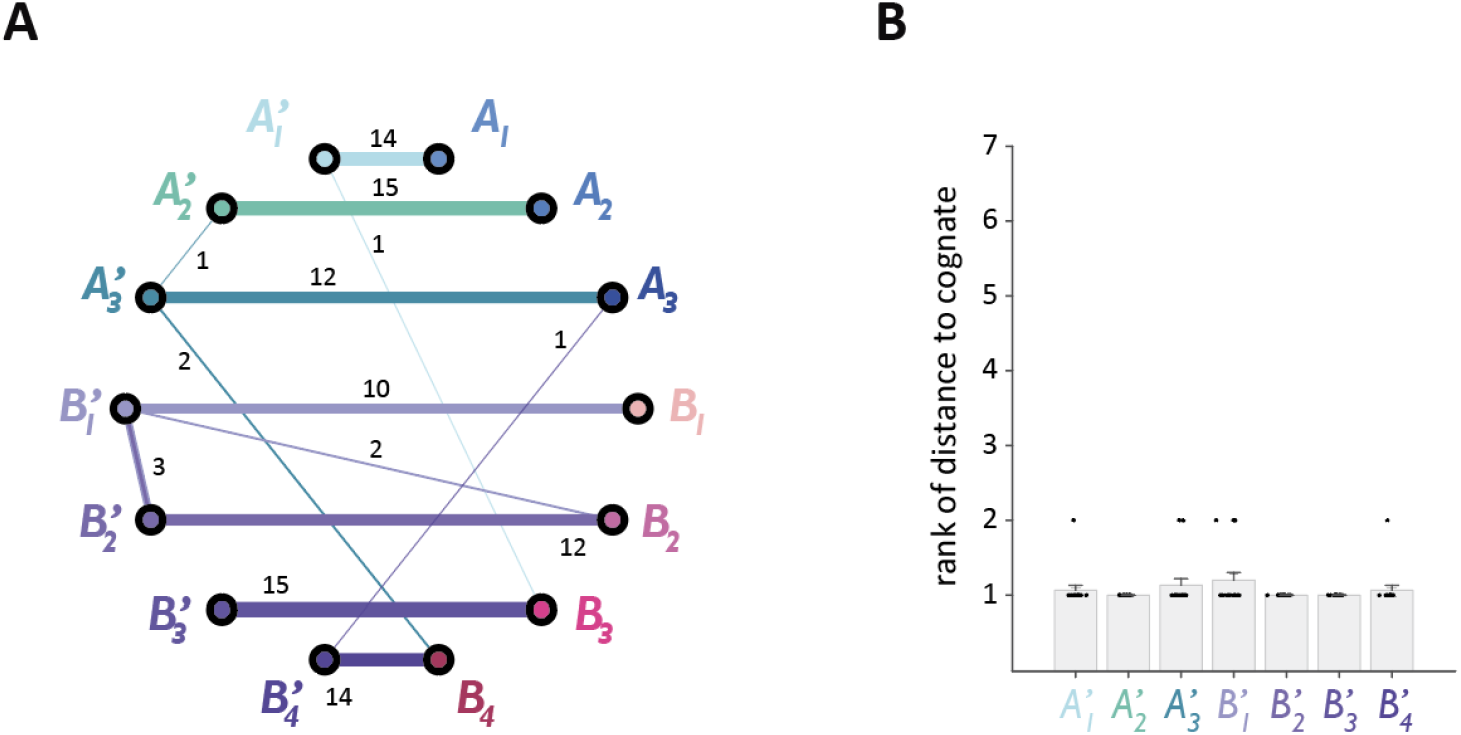
*A*’ and *B*’ neural activity clusters remain closest to their respective cognate clusters in *A* and *B*. **A**, Closest cluster distribution for each step in *A*’*B*’ across 15 sessions. In this analysis, the distances were calculated to all other 13 clusters (i.e. both within second order composite and to first order constituents. Nevertheless, in the majority of cases, the closest cluster is the congate one in the first-order constituent. **B**, Rank of the distance to the cognate cluster among all distances to just first-order constituent clusters for each step in *A*’*B*’. Dara are mean +/−S.E.M. n = 15 sessions. N = 5 animals.

**Figure S8.**
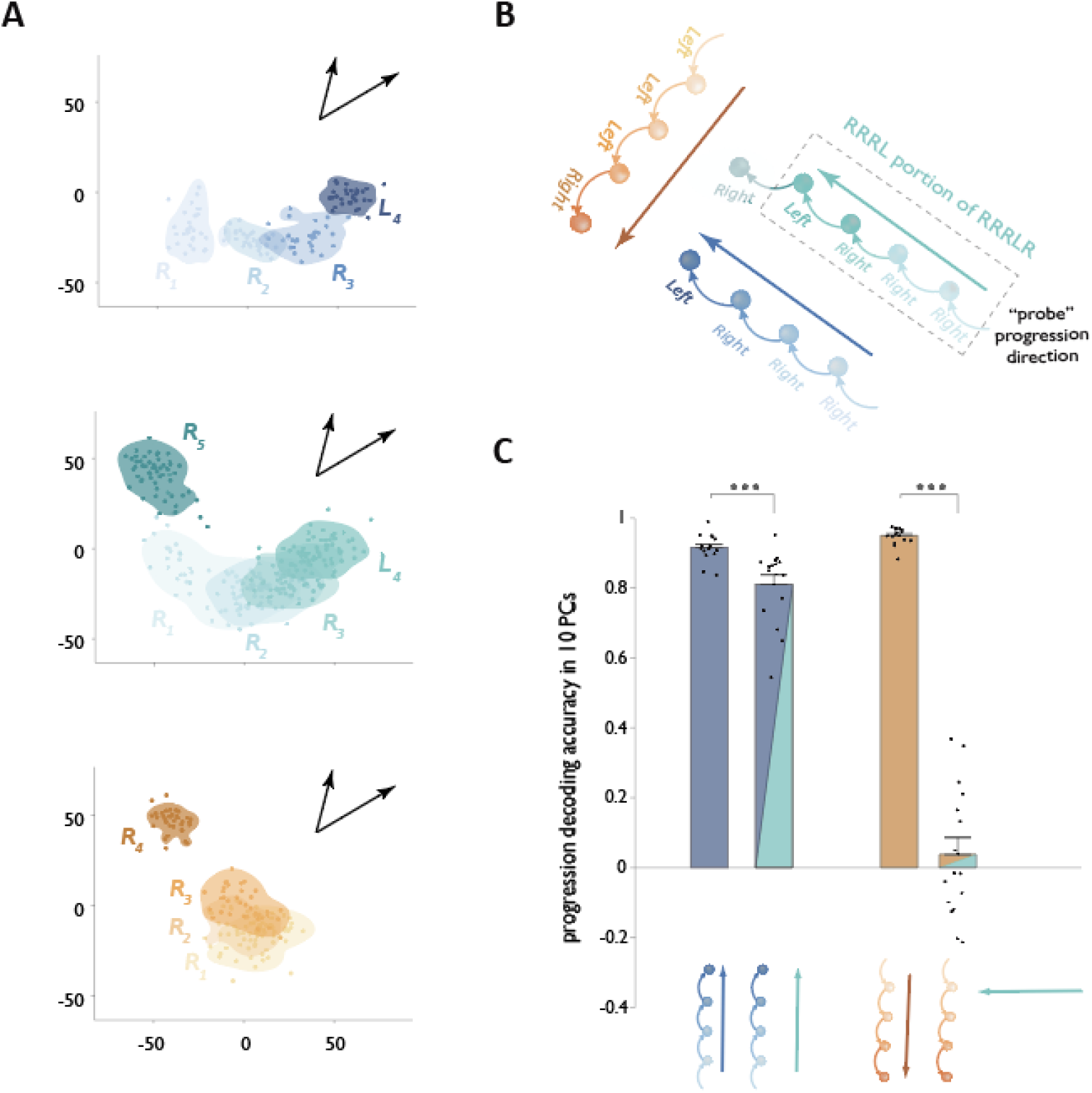
related to Fig. 4 | Representational geometry is similar for RRRL and RRRLR, but different from LLLR. **A**, Example neural geometry for *R*_1_*R*_2_*R*_3_*L*_4_ (top panel), for its one-letter extension, *R*_1_*R*_2_*R*_3_*L*_4_*R*_5_ (middle panel), as well as for the unrelated *L*_1_*L*_2_*L*_3_*R*_4_, plotted m the two dimensional space defined by the best *R*_1_*R*_2_*R*_3_*L*_4_ progression direction and the best residual _1_*R*_2_*R*_3_*L*_4_*R*_5_ progression direction (black arrows, see Methods). *R*_1_*R*_2_*R*_3_*L*_4_ and *R*_1_*R*_2_*R*_3_*L*_4_ progress along similar arcs. In contrast, no clrea progression axis for *LLLR* is evident in this projection plane. **B**, Schematic of the approach used to quantify alignment of the representations: RRRLR sequence was used to define a sequence-progression direction for its embedded *R*_1_*R*_2_*R*_3_*L*_4_ substructure. We then assessed how well this direction could generalize to capture sequence progression ifor *R*_1_*R*_2_*R*_3_*L*_4_ and *L*_1_*L*_2_*L*_3_*R*_4_, sequences. The underlying intuition is that if two sequences share an aligned structure in neural space, then the progression direction extracted from one should accurately decode progression through the steps of the other. Conversely, misaligned sequence representations would each yield poor decoding performance when using a direction derived from the other. **C**, Left bars: Progression decoding accuracy for *R*_1_*R*_2_*R*_3_*L*_4_ using either its native progression direction (blue arrow in B and C), or one determined by the *R*_1_*R*_2_*R*_3_*L*_4_ component of the *R*_1_*R*_2_*R*_3_*L*_4_*R*_5_ representation (cyan arrow in 8 and C). Note that while the decrease in the decoding accuracy is significant: two-tailed Wilcoxon signed-rank test: W = 135, p = 5.31 x 10^−^, n = 16, it is modest (cf. right bars). Right bars: Progression decoding accuracy for *L*_1_*L*_2_*L*_3_*R*_4_ using either its native progression direction (orange arrow) or one determined by the *R*_1_*R*_2_*R*_3_*L*_4_ component of the *R*_1_*R*_2_*R*_3_*L*_4_*R*_5_ representation (cyan arrow in B and C). Note the markedly worse decoding accuracy in the latter case. Two-tailed Wilcoxon signed-rank test: W = 136, p = 4.38 x 10^−4^, n = 16 Note a markedly more profound decrease in decoding accuracy (cf. left bars) One key implication of these results is that *L*_1_*L*_2_*L*_3_*R*_4_ and *R*_1_*R*_2_*R*_3_*L*_4_*R*_5_ have misaligned progression directions (as do their contractions, *L*_1_*L*_2_*R*_3_ and *R*_1_*R*_2_*L*_3_*R*_4_ respectively) Data are mean */-S.E.M.

**Figure S9.**
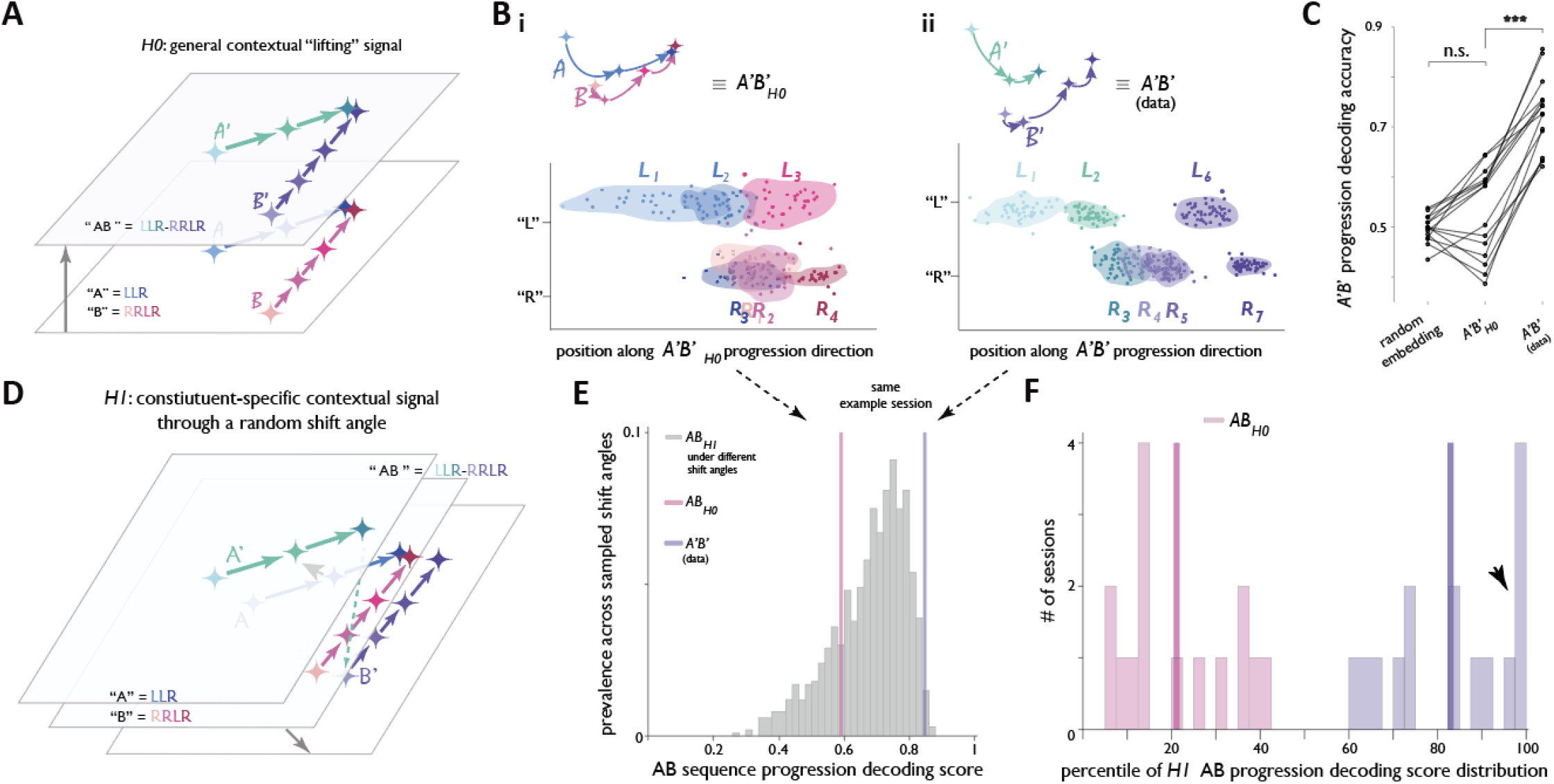
related to Figure 4 | Generic, randomly assigned contextual signal cannot explain representational geometry of AB composition. **A**, Transformation predicted under *H*_*o*_ assumes uniform contextual “lifting’’ of constituent representations such that the relative organization of *A*’ and *B*’ mirrors that of *A* and *B*. **B**, “Format” representation for an example behavioral session expected under *Ho* (i) and observed in data (ii). **C**, *A’B’* progression decoding accuracy expected under random embedding or *H*_*o*_, compared with the data. n=15 session, N=5 animals, wo-sided Wilcoxon signed-rank tests with Bonferroni correction (3 comparisons); random embedding vs A’B’_H0_: W-statistics=34, p=0.45; random embedding vs A’B’_(data)_: W-statistics=0, ps 1.8 ×10^−4^; A’B’_H0_, vs A’B’_(data)_: W-statistics=0, p = 1.8×10^−4^. **D**, Transformation predicted under *H*_*1*_assumes a common displacement of A-related clusters, and a common, but distinct displacement of B-related clusters. **E**, *A’B’* progression decoding scores for the example session in (B), shown relative to the distribution obtained under *H*_*1*_across 100 random shift-angle pairs (magenta, *H*_*o*_expectation: purple, data). **F**, Dataset-wide distributions of progression-decoding scores expected under *H*_*o*_or observed in data, expressed as percentiles within the corresponding *H*_*1*_distributions. Arrowhead denotes sessions falling outside the *H*_*1*_confidence interval.

**Figure S10.**
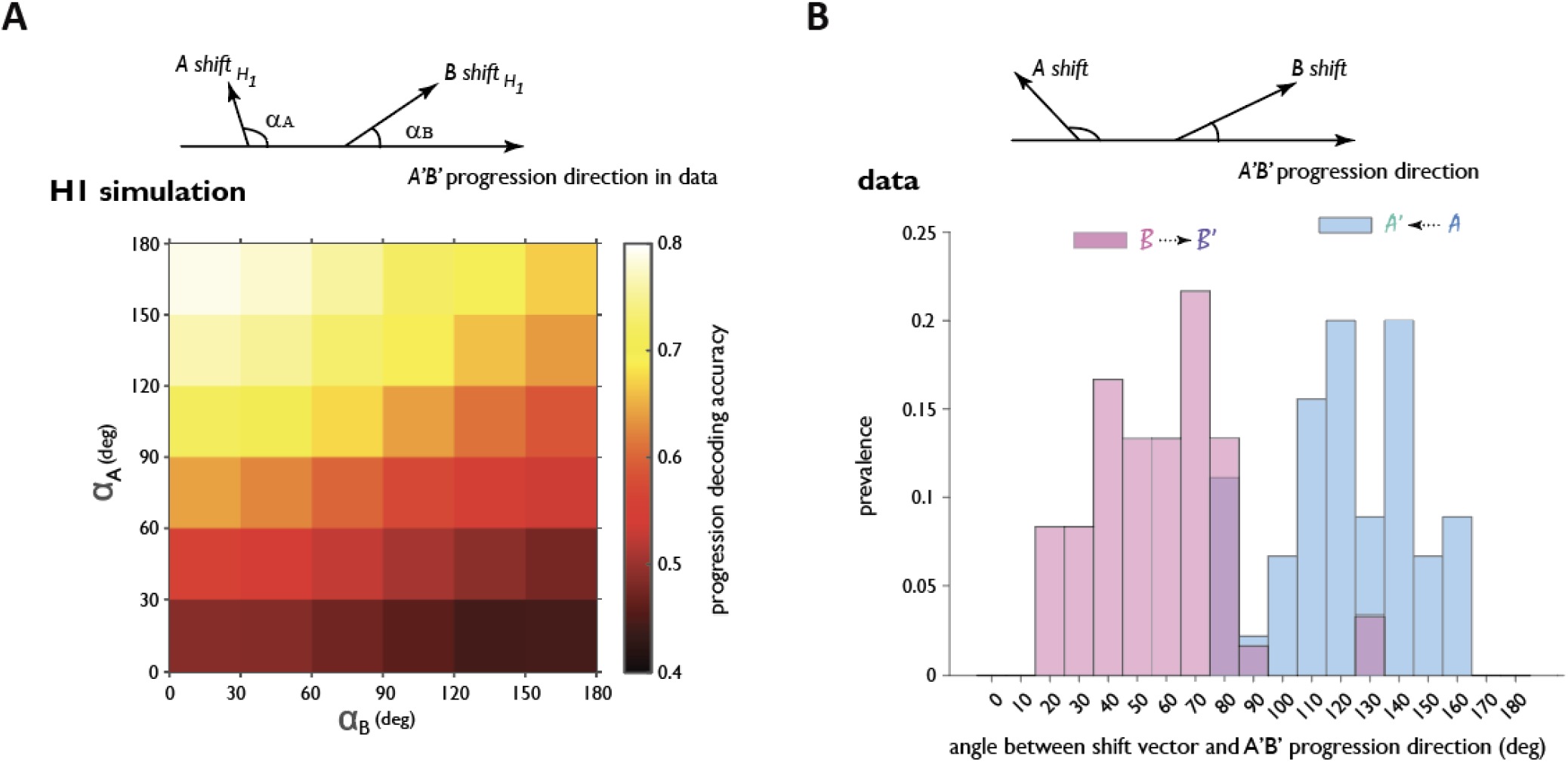
related to Figure 4 | *H*_*1*_shifts that push A related clusters earlier rnd B-related clusters later along the actual A’B’ progression direction achieve better progression decoding score. **A**, For *H*_*1*_simulation (random, independent A-shift and B-shift): correlation between progression decoding score and shift component to earlier (for A shift) and to later (for B-shift| along the empirical progression direction. **B**, Distribution in the data of angles between average A- or B-related cluster shifts and the A’B’ progression direction. The predominantly acute angles for B-shift is consistent with downward movement along A’B’, whereas obtuse angles for A-shift are consistent with upward movement.

**Figure S11.**
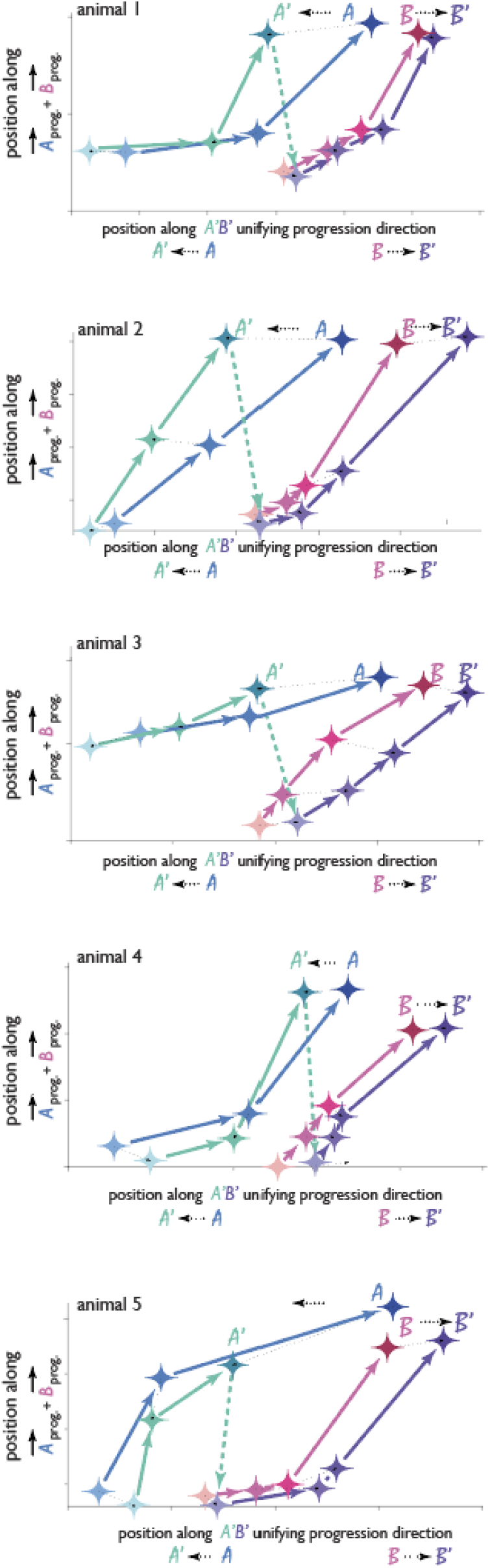
related to Figure 4 | Systematic shifts of *A*-clusters and *B*-clusters across animals. Geometry of *A*’, *B*’, *A* and *B* cluster centroids, projected onto the two-dimensional space defined by the *A’B’* progression direction and the average of *A* and *B* progression directions, shown for example sessions from five animals. A-related clusters are shifted upward and B-related clusters downward in the second-order composite.

**Figure S12.**
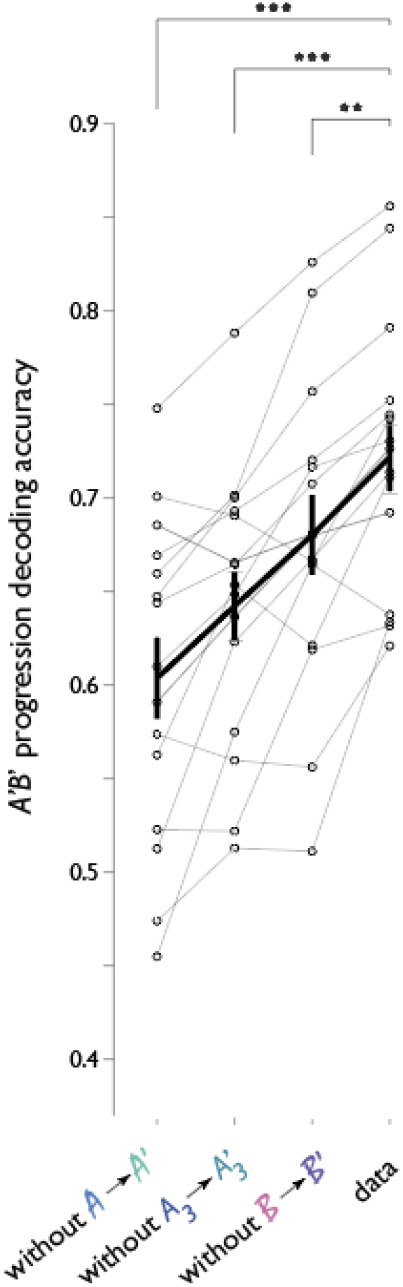
related to Figure 4 | Contributions to the establishment of format-adhereing unifying progression. *A’B’* progression decoding accuracy without *A* to *A’, B* to *B’* or without the unique displacement of *A*’_*3*_. n = 15 sessions, N=5 animals. Two-sided Wilcoxon signed-rank tests with Bonferroni correction: data vs without *A* to *A*; W=0, p=1.8×10^−4^; data vs without *A*_*3*_-> *A*’_*3*_, W=0, p=1.8×10^−4^; data vs without *B* to *B*’ W=4, p=1.3×10^−3^.

**Figure S13.**
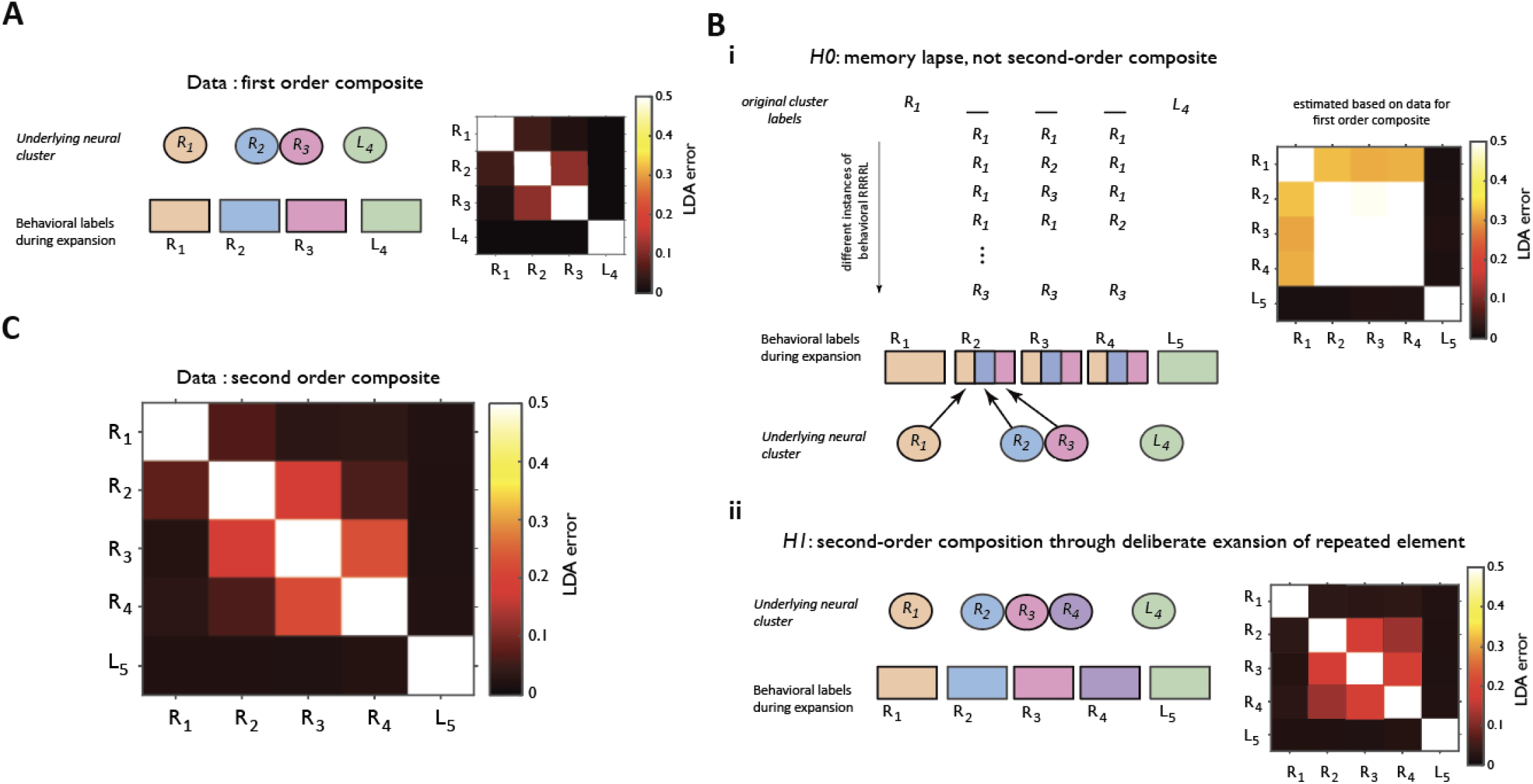
related to Figure 6 | Behavioral variants do not arise from a memory lapse. **A**, Confusion matrix for a linear classifier trained to distinguish neural activity associated with different sequence steps in the first-order composite RRRL. **B**, Expected confusion matrices for classifying behavioral steps in RRRRL expansions under two competing hypotheses. **i:** Under *H*_*o*_(memory lapse hypothesis), extra steps arise because the animal forgets which subsequent R it is executing, though it remembers the first step and final step (with a distinct motor action). When lapses occur, behavioral steps R2, R3, or R4 would randomly sample from the original *R1, R2* and *R3* neural activity clusters, predicting high classification confusion, **ii**, Under H, (compositional expansion hypothesis), each RRRRL step is represented by a distinct amFC neural activity cluster, predicting minimal classification error. **C**, Data are more consistent with *H*_*1*_arguing against memory lapse hypothesis. n = 9 sessions. N = 3 animals. Wilcoxon rank sum test for off-diagonal values for R2-R3-R4 vs 0.5 (in b ii): W = 0, p = 1.63 x 10^−9^, n = 48 See also Figure 6J (consistent positive displacement along progression direction) for additional evidence against a memory lapse.

**Figure S14.**
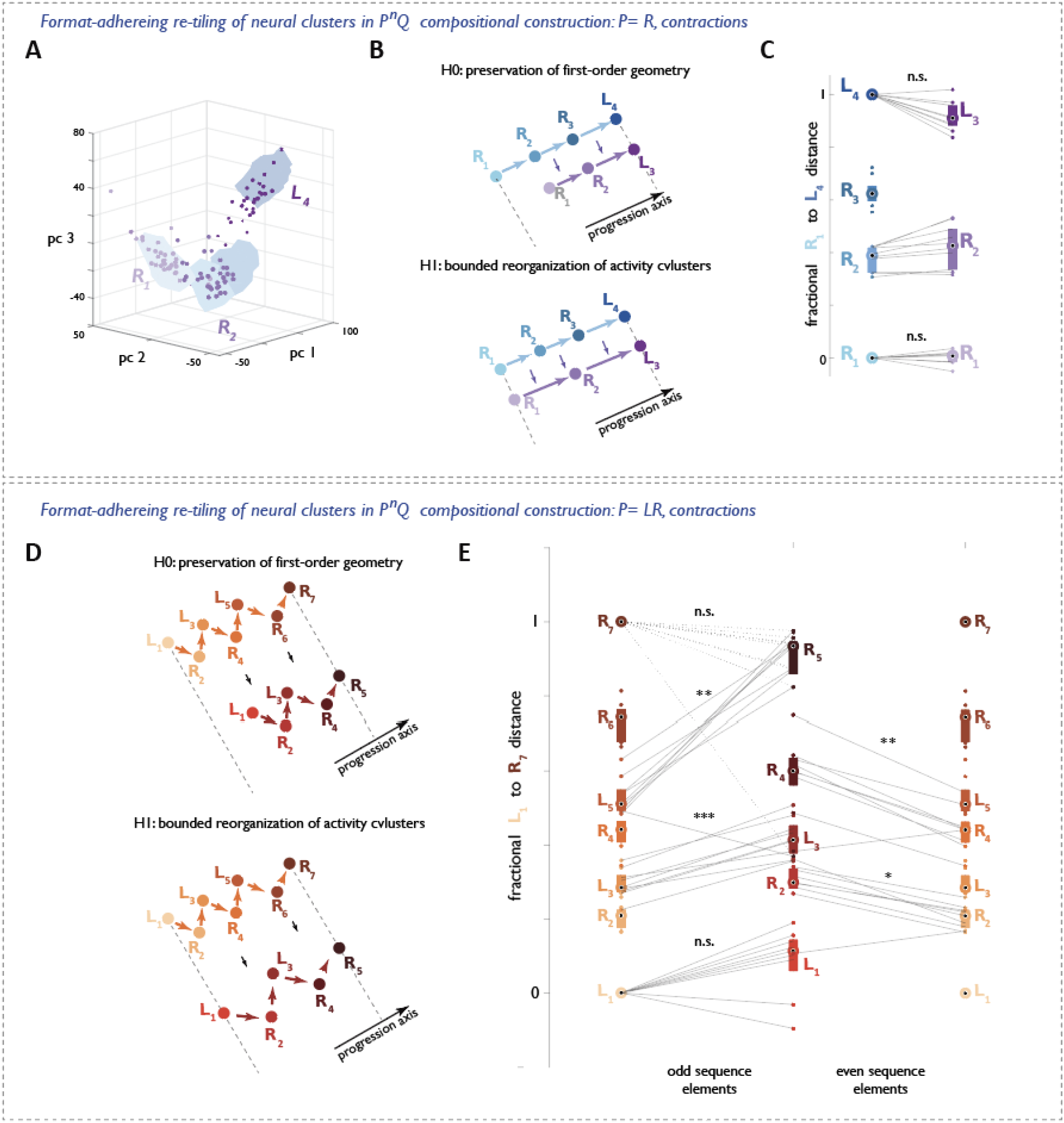
related to Figure 6 | Format-adhering re-tiling of neural clusters along the progression direction in P^n^Q compositional construction: contractions. **A**, Neural geometry for RRRL (top) and its compositional contraction RRL (bottom; *RRL* clusters overlaid on *RRRL* cluster outlines in blue) visualized in the space of the first three principal components for an example session. Points are colored by step identity. **B**, Schematic of the predicted representational transformation from RRRL to its contracted behavioral variant RRL under two hypotheses. Under *H*_*o*_, contractions preserves the first-order geometry, yielding consistent duster locations by removing the first clustrer (top). Under *H*_*1*_, expansion requires a rearrangement of cluster positions within the original boundary (bottom). **C**, Dataset-wide normalized positions of *R*_1_,*R*_2_,*R*_3_,*R*_4_and *R*_1_,*R*_2_,*L*_3_cluster centroids projected onto the *RRRL* progression axis. n=9 sessions, N=3 animals. **D-E** Same analyses as in (b–c), but for the transformation from LRLRLRR to its contracted variant LRLRR. n=9 sessions, N=3 animals. **, p<10^^^-2, ***, p<10^-3.

**Figure S15.**
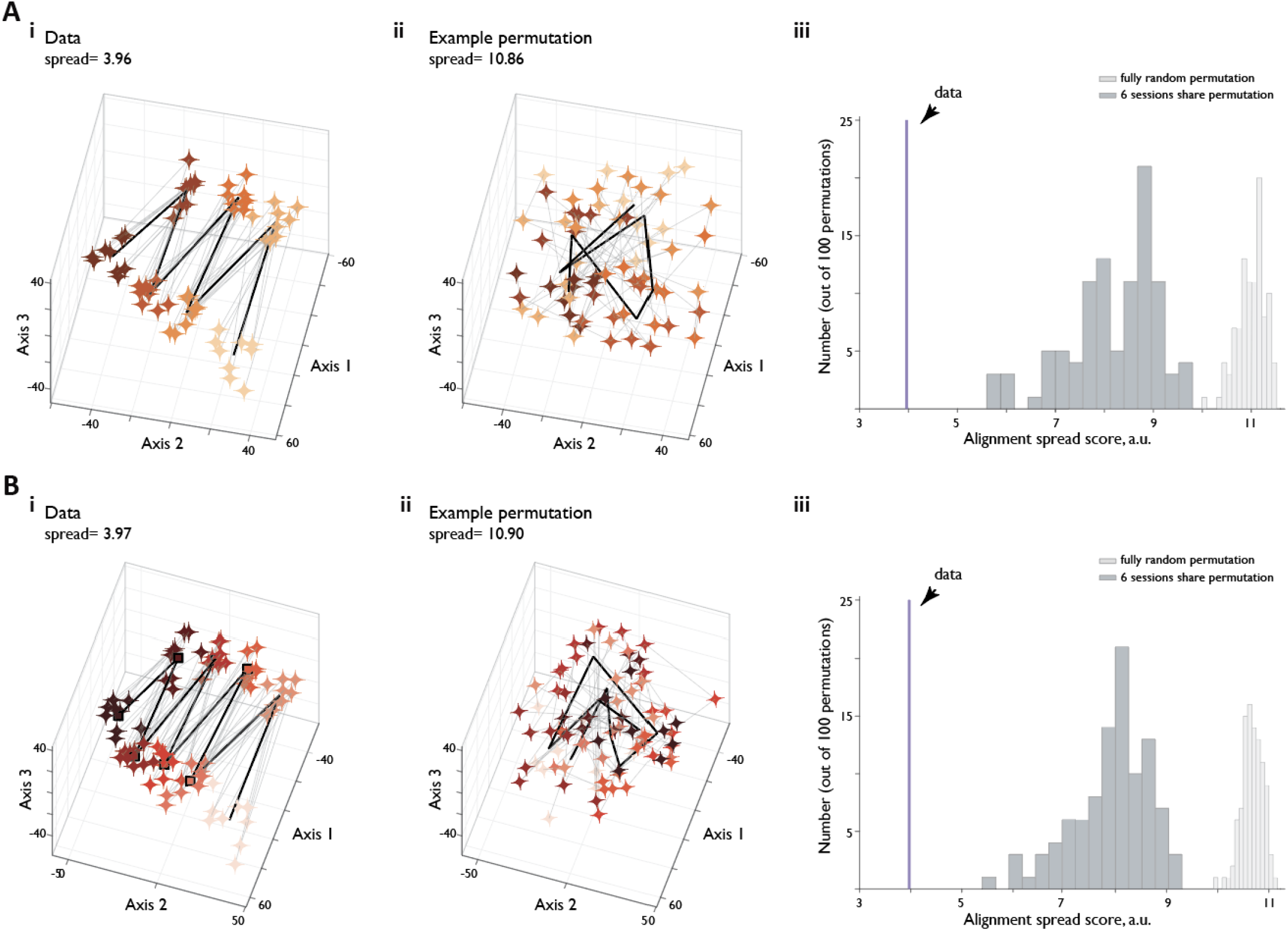
related to Figure 7 | Representational alignment in P^n^Q composition is not a consequence of random emnedding in a high dimensional space. **A, i:** Global alignment of (*LR*)^*3*^*R* shapes across animals and sessions, **ii:** Alignment after a single random permutation of ordinal labels within each session, **iii:** Distribution of spread (quality of alignemnt) scores across 100 fully random ordinal label permutations, and across 100 permutations where half of the sessions shared the permutation structure. The spread score observed in the data falls far outside of both distributions. p<0.01 by design (100 permutations). **B**, Same as in (a) but for (*LR*)^*4*^*R* shapes. n=9 sessions, N=3 animals

